# Field persistence of entomopathogenic fungi as biocontrol agents and network-level associations within the soil microbial community

**DOI:** 10.1101/2025.11.02.686131

**Authors:** Sabina Matveev, Victoria Reingold, Adi Faigenboim, Yaara Livne, Michael Davidovitz, Dana Ment

## Abstract

The effects of entomopathogenic fungi (EPF) on native soil microbiota are poorly understood. Here, we investigated the persistence of the EPF *Metarhizium brunneum* and *Beauveria bassiana* as preventive treatments for date palms against the red palm weevil, *Rhynchophorus ferrugineus*, and monitored the microbial community within the palms. We assessed the changes in the soil microbiota of the palms following application by fungal soil isolation techniques, high-throughput metagenomics, EPF persistence assays, and accompanied that with a continuous monitoring of date palm health. EPF preemptive treatments significantly improved palm protection against the weevil. While both EPF species persisted for 180 days post-application, their effectiveness diverged, with *B. bassiana* providing longer palm protection. Overall microbial community composition showed no major changes following EPF applications; however, transient alterations increased the relative abundance of local EPF genera, with dominant taxa being more likely to persist. Both EPF treatments enhanced the relative abundance of nitrogen-fixing bacteria, organic matter decomposers, and ammonia-oxidizing archaea, suggesting potential implications for nutrient cycling that require further validation through direct functional measurements. *B. bassiana* induced the strongest temporal effects on diversity indices, while *M. brunneum* was more persistent but showed no significant changes in diversity indices compared to the control. These results highlight EPF as a multifunctional biocontrol agent – moderately shaping soil microbiomes for beneficial ecosystem functions.

## 1. Introduction

The soil microbiome shapes plant health by driving nutrient cycling, disease resistance, and stress tolerance^1,2^. While rhizobia, mycorrhizal fungi, and biocontrol agents, such as entomopathogenic fungi (EPF), are well-studied, the effectiveness of EPF in agriculture is inconsistent, and their interactions with native microbiota remain unclear^3–9^. Traditional methods (e.g., T-RFLP) provided limited resolution ^6^, while high-throughput sequencing enables finer taxonomic and functional insights. Previous studies (e.g., Canfora^6^, Hirsch^10^) have shown that *Beauveria bassiana* can persist in soils, albeit with minimal long-term community shifts, underscoring the need for viability assays and high-resolution sequencing. EPF are of interest due to their role as natural pathogens of arthropods and insects. EPF are common components of the soil microbiota^11^, and their presence is positively correlated with factors such as organic matter content^12^. The diversity and abundance of EPF in soils are influenced by various environmental factors, including abiotic factors, such as temperature, humidity, radiation, and soil properties, as well as biotic factors, including competition, antagonism, and symbiosis with other microorganisms^11–13^. Their high abundance and diversity across ecosystems and soil types make them important factors in integrated pest management (IPM) and soil fertility^12,14^.

Here, we asked what taxonomical and functional changes occur in the soil microbiota following EPF application in *Phoenix dactylifera* plantation infested by the red palm weevil (RPW), *Rhynchophorus ferrugineus* (Coleoptera: Curculionidae). We hypothesized that the inundative application of EPF as a myco-biopesticide in date palm plantations would enhance their persistence in the soil, and potentially alter the abundance and diversity of the soil microbial community while conferring protection to the palms against the RPW. The mode of EPF conidia dissemination by RPW adults toward eggs and larvae following inundative application on surfaces was demonstrated by us previously in a simulation system^16^. Furthermore, we aimed to evaluate the effect of EPF treatment under field conditions and changes in palm health and the initial microbial composition of the soil, which may play a role in the fungi’s long-term persistence. The specific objectives of this study were to: measure the persistence of the EPF over time, correlate soil microorganism diversity with EPF persistence to identify the associated key taxa; characterize the soil microbial community around palms in the Jordan Rift Valley following application of EPF; and study the principal metabolic functions and diversification of soil microorganisms in the presence of EPF. All objectives were conducted within the overarching framework of integrating EPF into IPM strategies against RPW.

## 2. Results

### 2.1. Velifer and Mb7 Treatments Protect Palm while Exhibiting High Soil Persistence

Tree health was monitored using sensors as described previously^1^, on 15 trees sampled at 5 time points (pre-application, application day, and 30, 90, and 180 days post-application) starting in January 2019. Each treatment [Control, Mb (*Metarhizium brunneum* strain Mb7), and Velifer (*Beauveria bassiana*)] was applied to five palm trees randomly positioned within the plot. A single tree (no. 73) was excluded due to an infestation report at the onset of the experimental period. A significant effect of EPF treatment and time was observed (Ordinary least squares (OLS) regression; F = 5.580, p < 0.00009), explaining 14.7% of the variance in health rates (R² = 0.147, adj. R² = 0.121). Baseline health rate was 78.09% (p < 0.0001), with overall decline across treatments (days coefficient = –0.0868, p < 0.00022). Interaction terms revealed contrasting trends in treatment efficacy: Mb decreased over time (coefficient = – 0.1311, p = 0.002), Velifer increased (0.0951, p = 0.042), and control was non-significant (–0.0508, p = 0.236) (Fig. 1c). Model diagnostics (AIC = 1577.3, BIC = 1596.1) confirmed robustness **(Fig. 1c; Supplementary Tables 1–4)**.

**Fig. 1.**
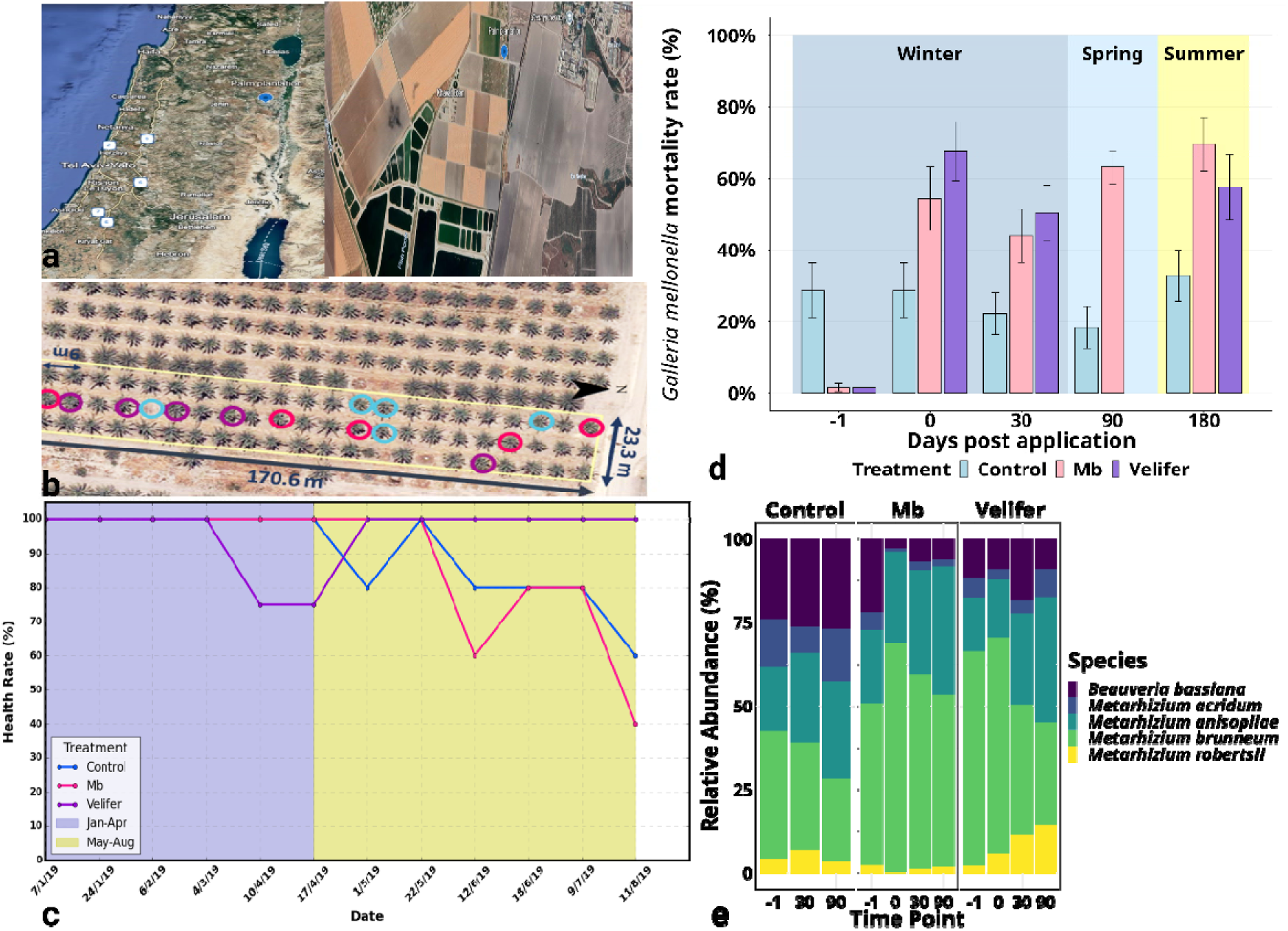
Experimental layout and EPF persistence in the soil. **a** Map of the research site (Eden Farm, Beit-She’an Valley, Israel). **b** Plot layout (23.3 m × 170.6 m) with treatment subdivisions, blue-control; pink-Mb, and purple-Velifer; circles indicate sampling points. **c** Tree health rates over time for each treatment (control, Mb [*Metarhizium brunneum* strain Mb7] and Velifer [*Beauveria bassiana*]), shown across two seasonal periods: January–April (purple shading) and May–August (yellow shading). Treatments were applied on January 7, 2019, with soil samples taken before, immediately after, and at 30- and 90-day post-application intervals (Supplementary Table 5). **d** EPF infectivity measured via *Galleria mellonella* baiting on soil samples through the 180-day period, Velifer 90-day data were missing due to technical issues (Supplementary Table 6). **e** Relative EPF abundance in soil was visualized in stacked bar plots across treatments and time points (–1, 0, 30, 90 days), with control lacking time 0 as a negative control and Velifer missing day 90 due to technical issues.

EPF persistence was examined by *G. mellonella* larvae mortality rate in soil and bark samples across 4 time points, 3 treatments (Control, Mb and Velifer) based on random sampling location around each palm tree. The effects of time (F(4,4) = 14.48, p < 0.0001), treatment (F(2,2) = 21.82, p < 0.0001), and sample type (F(1,1) = 32.29, p < 0.0001) on fungal efficacy were statistically significant by ANOVA. **(Fig. 1d; Supplementary Table 6)**. Soil samples had higher mean mortality rates than bark samples in combined treatments analysis (40.2% and 25.3, respectively; *p*<0.0001). Mortality rate was highest on the day of application and on day 180, and lowest at day −1 in combined treatments analysis (43.6, 37.9, and 14.5, respectively; *p*<0.0001). Mb and Velifer showed similar mortality rates, both significantly higher than in control rates, in a combined time points analysis (39.5 and 38.3, respectively; *p*<0.0001).

### 2.2. Trends of EPF within each treatment over time in soil

To understand the dynamics underlying these palm protection patterns, we examined treatment-specific temporal trends in EPF composition. Before first EPF application (time: –1), soil around control trees showed significantly (Student’s t: t=1.993, p<0.001) higher mortality in Galleria-bait assays (28.8%) despite balanced fungal populations, compared to treated trees produced very low mortality (1.5–1.6%) (Fig. 1d). At application (0), mortality sharply increased in the EPF treated trees (Mb 54.4%, Velifer 67.7%), but remained stable in control (28.8%) (*p<*0.001). From 30– 180 days, mortality remained higher in treated trees, Mb increased from 44 to 69.6%, and Velifer to 63.2%, while control maintained lower mortality **(Supplementary Table 7, Fig. 1d)**. These results were also supported by the relative abundance in species-level **(Fig. 1e)**. In soils surrounding control trees, we found naturally occurring EPF with dynamic shifts *B. bassiana* increased over time*, M. acridum* peaked at 90 days, *M. anisopliae* rose steadily to 90 days, *M. brunneum* peaked at 30 days and *M. robertsii* rose at 30 days and then declined. In Mb treated trees, *B. bassiana* was stable with rises at 30 and 90 days, *M. anisopliae* increased at 90 days, *M. brunneum* decreased at 30 and 90 days, and *M. robertsii* remained low, and *M. anisopliae* and *M. brunneum* dominated by 90 days. In Velifer-treated trees, *M. acridum* peaked strongly at 90 days*, M. anisopliae* increased gradually to 90 days, *M. brunneum* increased moderately (less than in Mb), and *M. robertsii* increased at 90 days **(Fig. 1e)**.

### 2.3. Temporal microbial community shifts and functional impacts of EPF treatments

To test whether treatments affect microbial composition or if other factors play a dominant role, we conducted a Principal Components Analysis (PCA). As we observed no clear separation by treatment parameter (Fig. 2a), we used tree location within the plot for the second analysis. The spatial tree location resulted in 3 distinct groups in the PCA (Fig. 2b). The results indicate that the applied treatments had no significant effect on overall microbial composition over time, with intra-plot variance occurring within the plot itself (Fig. 2c).

**Fig. 2.**
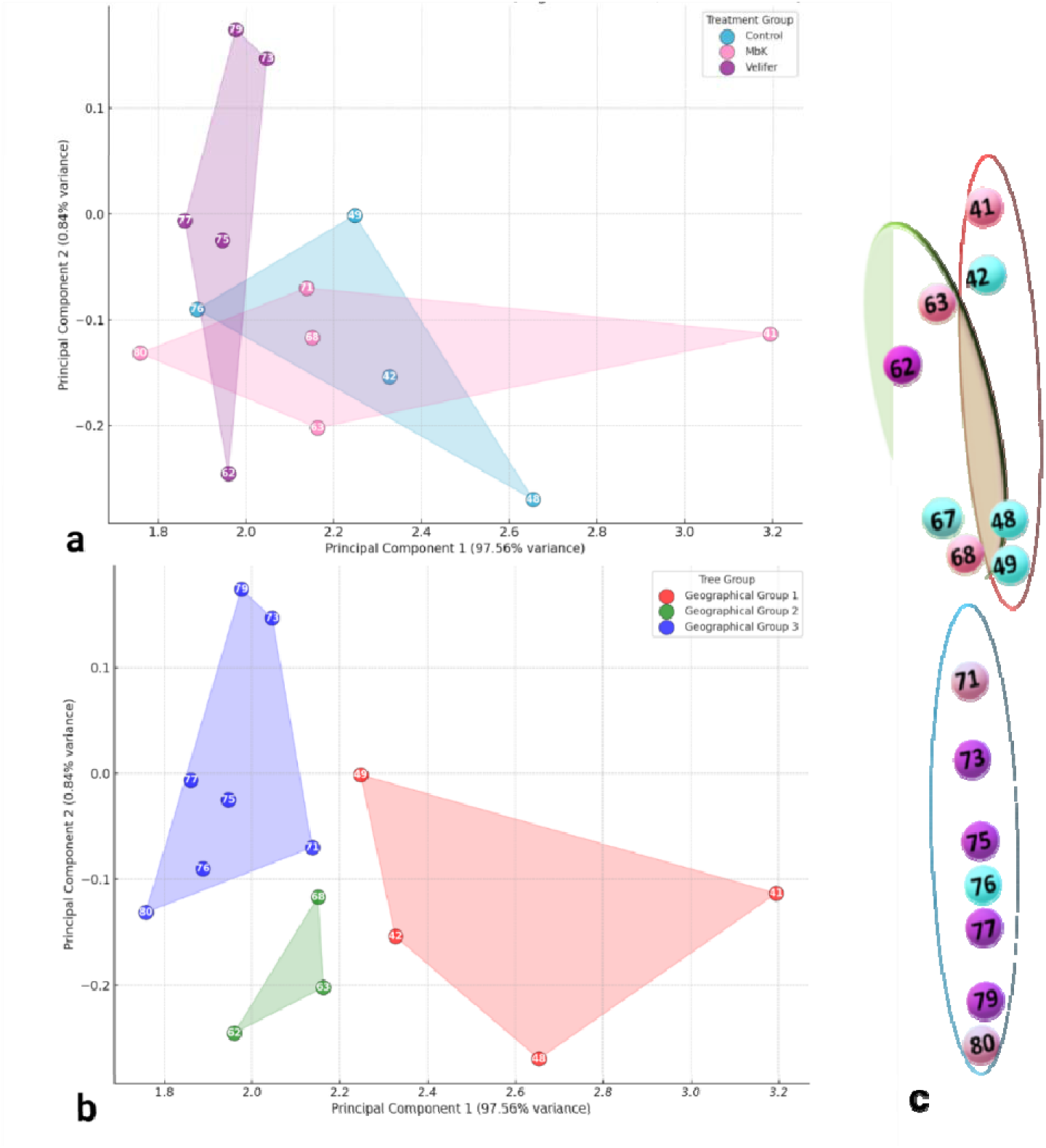
Principal component analysis (PCA) based on microbial relative abundance showing the distribution of the average tree samples. PCA was generated based on **a** treatment (control, Velifer and Mb) and **b** trees location, **c** samples spatial map.

Temporal shifts in microbial communities demonstrated by non-metric multidimensional scaling (NMDS) ordination (stress: 0.115–0.167, all <0.2) with PERMANOVA (Fig. 3a). The pre-application (–1) showed high stress (0.167) and overlapping groups. At time 0, treatments separated slightly (R² = 0.072, *p* value = 0.169; stress = 0.136). Maximum differentiation occurred at day 30 (R² = 0.176, *p* value = 0.002; stress = 0.115), with clear treatment separation, which persisted up to day 90 (R² = 0.098, *p* value = 0.023; stress = 0.162), indicating significant, time-dependent effects lasting at least 90 days.

**Fig. 3.**
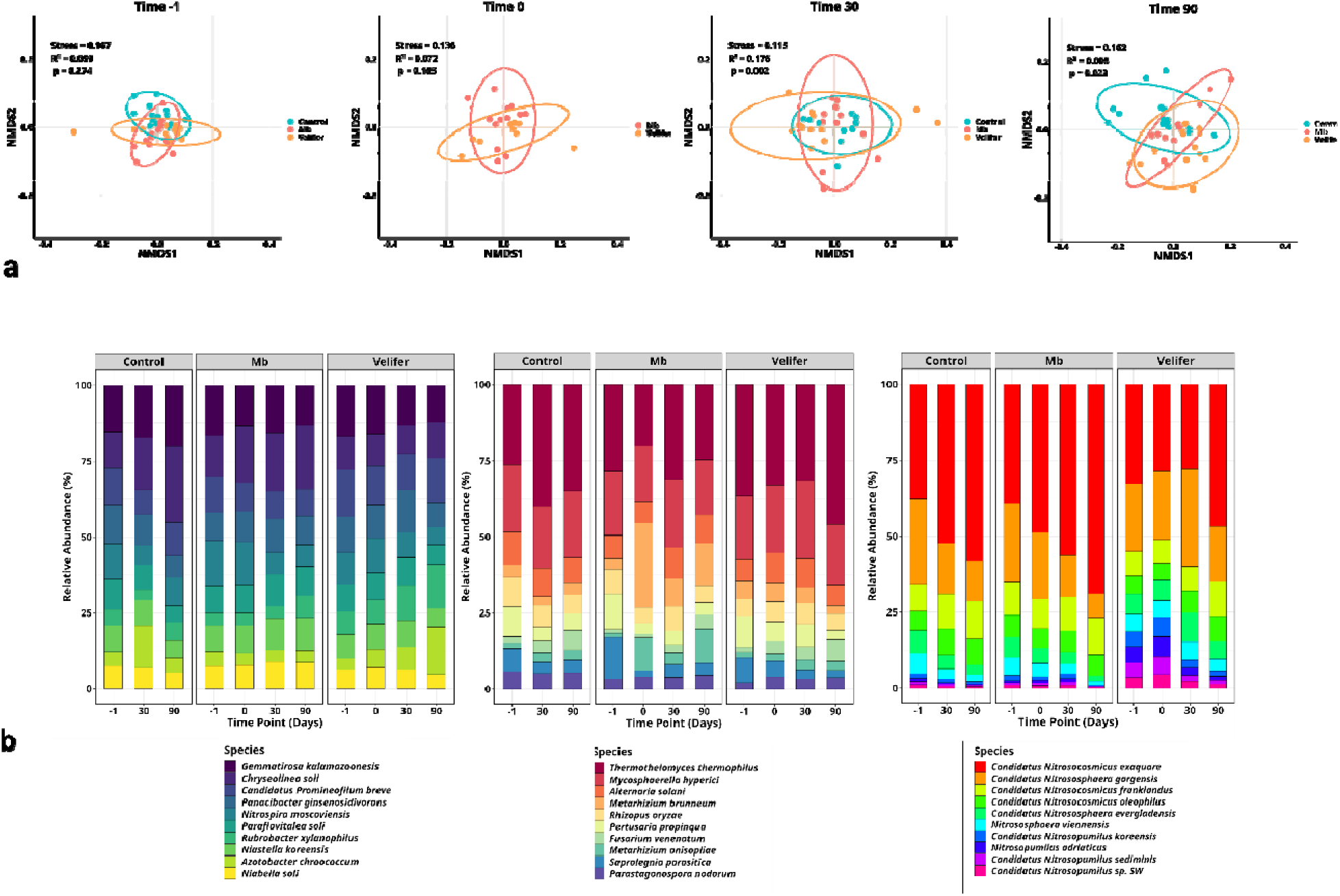
Microbial composition of treatment groups. **a** non-metric multidimensional scaling ordination of microbial communities with 95% confidence ellipses for each treatment (control; light blue, Mb; pink, Velifer; orange) at –1, 0, 30, and 90 days; PERMANOVA p-values, stress and correlation co-efficient shown on plots. **b** Relative abundance stacked bar plots of the top 10 most abundant bacterial (1), fungal (2), and archaeal (3) species for each treatment and time point.

**Figure 1.**
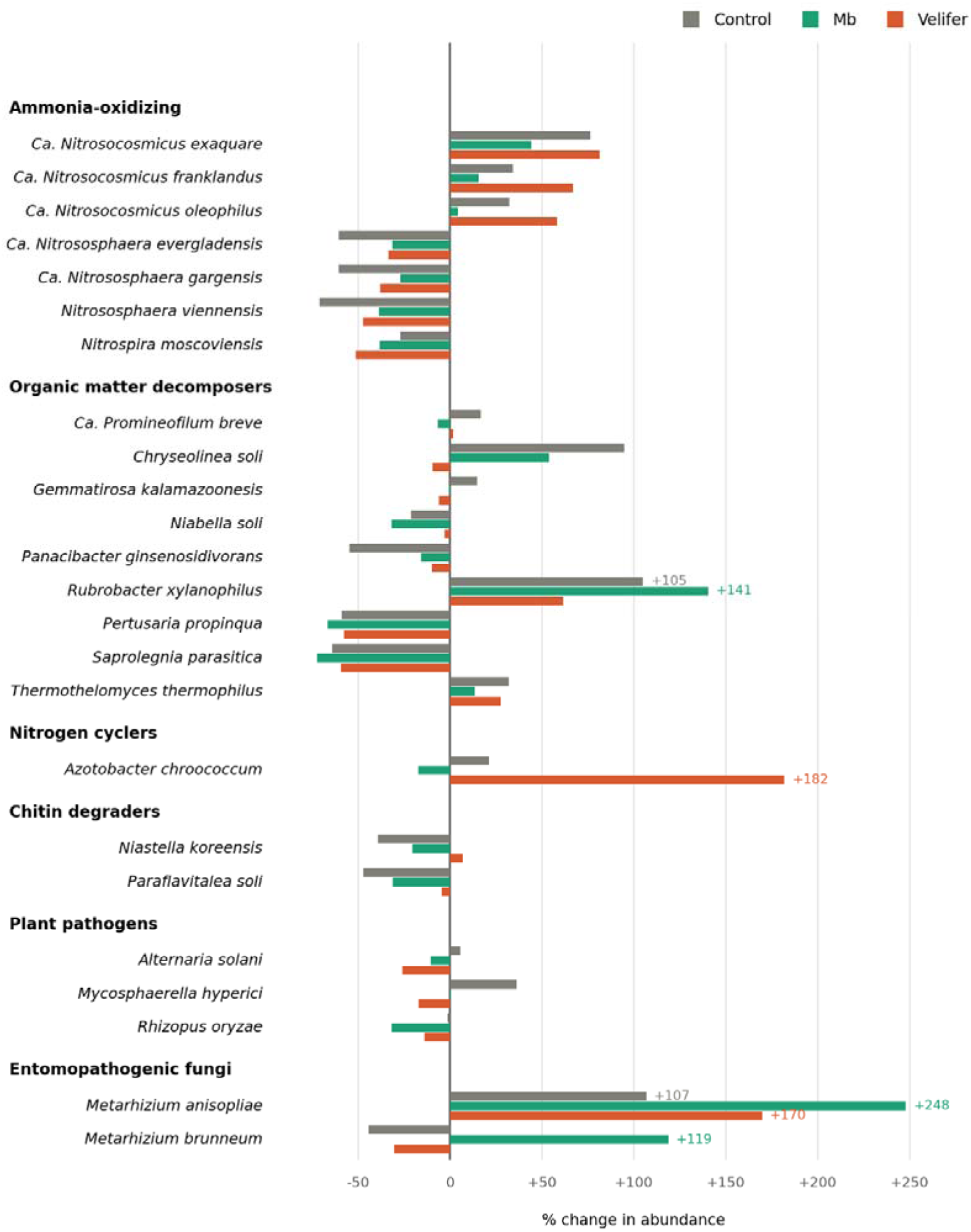
Treatment-induced changes in the relative abundance of soil microorganisms, grouped by functional guild. Heatmap showing the percent change in relative abundance (Δ%) of 24 microbial taxa following each of three treatments — Control, Mb, and Velifer [Mb *Metarhizium brunneum*– and *Beauveria bassiana*–based biopesticides, respectively] — relative to [baseline/untreated soil] at [time point; n = X per treatment]. Rows are individual taxa organised by functional group (ammonia-oxidizing, organic-matter decomposers, nitrogen cyclers, chitin degraders, plant pathogens, and entomopathogenic fungi); columns are the three treatments. Cell color encodes the direction and magnitude of change, from a decrease (blue) through no change (white) to an increase (red), with the exact value printed in each cell. To preserve visual contrast among the more moderate responses, the colour scale is clamped at ±120%; values exceeding this range (e.g. *Metarhizium anisopliae*, +248.1%) are shown at the scale extreme but retain their true value in-cell. Coloured squares beside each taxon name denote domain (purple, Archaea; blue, Bacteria; orange, Fungi). “Ca.” denotes *Candidatus*. [State statistical test and significance threshold; mark significant changes if applicable.]

**Figure 2.**
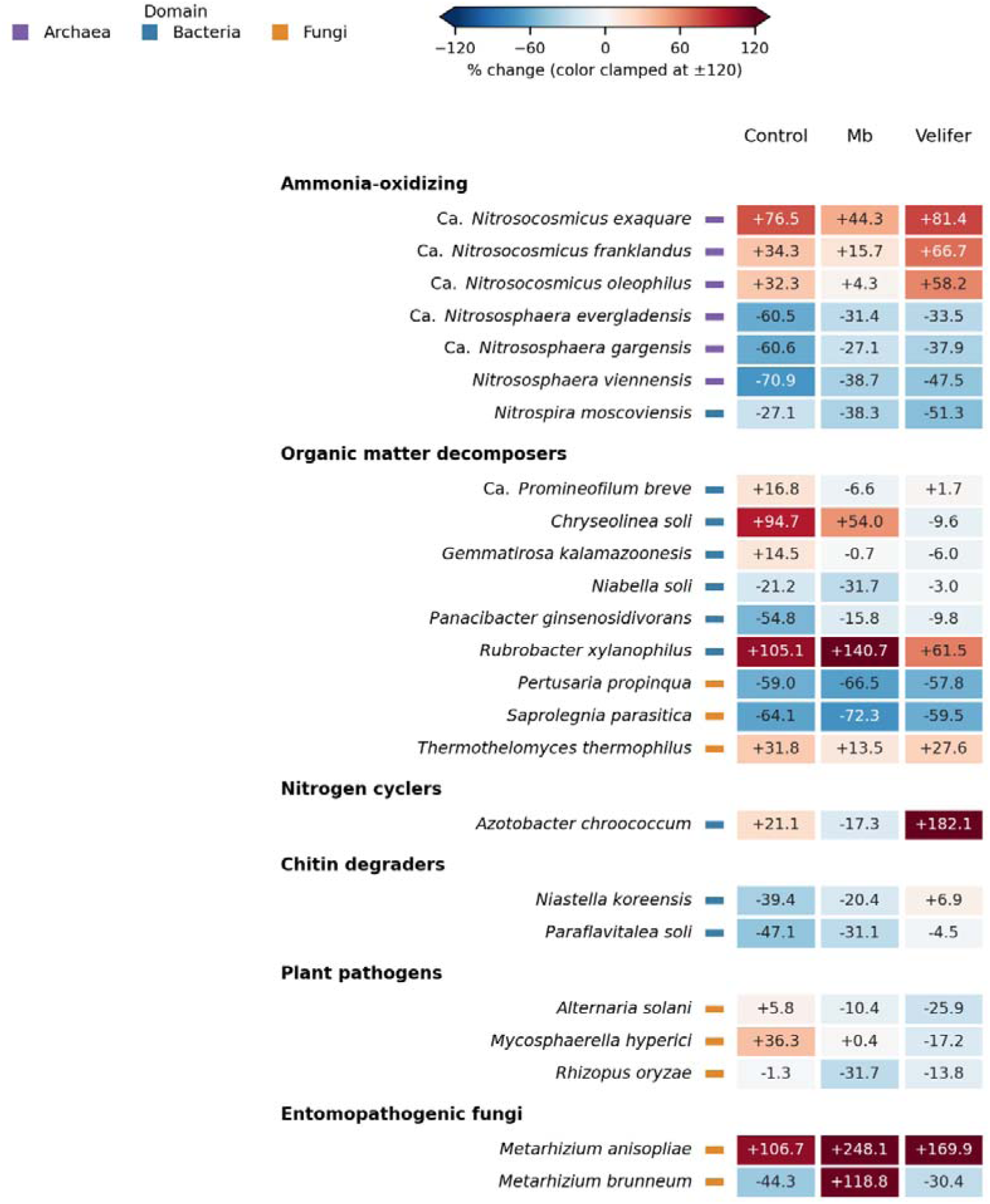
Direction and magnitude of treatment-induced abundance changes across functional guilds. Diverging horizontal bar chart of the percent change in relative abundance (Δ%) of the same 24 microbial taxa under the three treatments — Control (grey), Mb (green), and Velifer (orange) — relative to [baseline] at [time point; n = X per treatment]. For each taxon, the three bars represent the three treatments; bars extend leftward for a decrease and rightward for an increase from the zero reference line. Taxa are grouped by functional guild as in Figure 1. Unlike the heatmap, the axis is unclamped to show the full magnitude of response; the five responses exceeding +100% are labelled with their values. “Ca.” denotes *Candidatus*. [State statistical test and significance threshold; indicate significant responses, e.g. with asterisks, if applicable.]

**Table 1.** Microorganisms’ relative abundance changes due to treatment between pre-treatment and 90 days post-application, sorted by ecological function.

| Functional group | Microorganism | Domain | Control | Mb | Velifer |
| --- | --- | --- | --- | --- | --- |
| Ammonia-oxidizing | *Candidatus Nitrosocosmicus exaquare* | Archaea | 76.5 | 44.3 | 81.4 |
|  | *Candidatus Nitrosocosmicus franklandus* | Archaea | 34.3 | 15.7 | 66.7 |
|  | *Candidatus Nitrosocosmicus oleophilus* | Archaea | 32.3 | 4.3 | 58.2 |
|  | *Candidatus Nitrososphaera evergladensis* | Archaea | -60.5 | -31.4 | -33.5 |
|  | *Candidatus Nitrososphaera gargensis* | Archaea | -60.6 | -27.1 | -37.9 |
|  | *Nitrososphaera viennensis* | Archaea | -70.9 | -38.7 | -47.5 |
|  | *Nitrospira moscoviensis* | Bacteria | -27.1 | -38.3 | -51.3 |
| Organic matter decomposers | *Candidatus Promineofilum breve* | Bacteria | 16.8 | -6.6 | 1.7 |
|  | *Chryseolinea soli* | Bacteria | 94.7 | 54 | -9.6 |
|  | *Gemmatirosa kalamazoonesis* | Bacteria | 14.5 | -0.7 | -6 |
|  | *Niabella soli* | Bacteria | -21.2 | -31.7 | -3 |
|  | *Panacibacter ginsenosidivorans* | Bacteria | -54.8 | -15.8 | -9.8 |
|  | *Rubrobacter xylanophilus* | Bacteria | 105.1 | 140.7 | 61.5 |
|  | *Pertusaria propinqua* | Fungi | -59 | -66.5 | -57.8 |
|  | *Saprolegnia parasitica* | Fungi | -64.1 | -72.3 | -59.5 |
|  | *Thermothelomyces thermophilus* | Fungi | 31.8 | 13.5 | 27.6 |
| Nitrogen cyclers | *Azotobacter chroococcum* | Bacteria | 21.1 | -17.3 | 182.1 |
| Chitin degraders | *Niastella koreensis* | Bacteria | -39.4 | -20.4 | 6.9 |
|  | *Paraflavitalea soli* | Bacteria | -47.1 | -31.1 | -4.5 |
| Plant pathogens | *Alternaria solani* | Fungi | 5.8 | -10.4 | -25.9 |
|  | *Mycosphaerella hyperici* | Fungi | 36.3 | 0.4 | -17.2 |
|  | *Rhizopus oryzae* | Fungi | -1.3 | -31.7 | -13.8 |
| Entomopathogenic fungi | *Metarhizium anisopliae* | Fungi | 106.7 | 248.1 | 169.9 |
|  | *Metarhizium brunneum* | Fungi | -44.3 | 118.8 | -30.4 |

### 2.4. Temporal treatment effects on microbial diversity

Pairwise comparisons of Shannon diversity were performed between treatments. To evaluate whether Shannon diversity differed among the three treatments (control, Mb, Velifer), a one-way ANOVA was conducted with treatment as the single fixed factor (F[2, 136] = 4.27, p = 0.016), followed by Bonferroni-corrected pairwise t-tests between treatment pairs. The Shannon diversity index for the Velifer treatment was significantly lower than that for both the control (padj = 0.034) and Mb (padj = 0.0432) treatments. In contrast, control and Mb treatment did not differ significantly (p_adj_ = 1) (Fig. 4a, Supplementary Table 11a). The difference in Shannon indices between day −1 (6.783 ± 0.071) and day 90 (6.590 ± 0.282) showed the highest significance (t = 4.25, df = 44.97, FDR = 6.36E-04), suggesting a decrease in Shannon diversity.

**Fig. 4.**
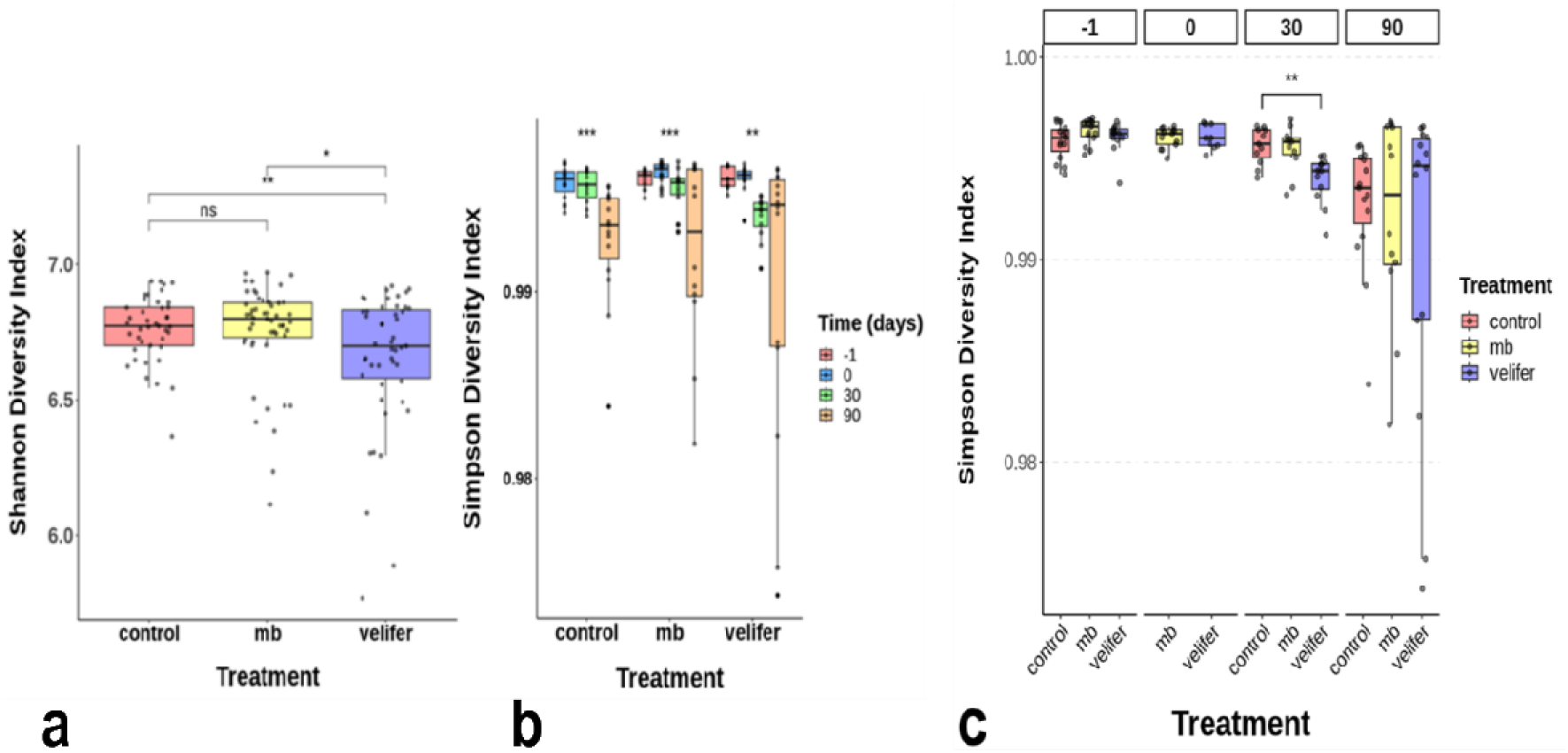
Alpha diversity indices across treatments and time points. Welch t-tests were performed for each pair of time points. Shannon diversity (base 10) was calculated with Benjamini–Hochberg FDR adjustment. **a** Shannon diversity across treatments (control, Mb, Velifer). **b** Simpson diversity across time (–1, 0, 30, 90 days) within each treatment. **c** Simpson diversity between treatments at each time point.

To select the most extreme effect over time, Welch’s t-test was performed for −1 vs. 30/90 days and 0 vs. 30/90 days. Both comparisons gave significant *p-values*. As we aimed to compare the most extreme situations in the plot following EPF application, we analyzed the datasets for day −1 and day 90 post-application for each treatment, thereby assessing the principal changes that occurred over time and treatments (t = 4.25, FDR = 6.36E-04, *p* < 0.001) **(Supplementary Table 11b)**. Shannon diversity analysis revealed a reduction between treatments (Fig. 4a), temporal effects within each treatment (Fig. 4b), and temporal effects between treatments (Fig. 4c). This analysis yielded a significant Simpson diversity index between control and Velifer at 30 days post-application. Significant changes in the Simpson diversity index indicated a temporal interaction effect within each treatment (control p = 0.000252, Mb p = 0.000295, Velifer p = 0.00791) and a significant dominance effect 30 days post-application (p = 0.0012) induced by Velifer compared with the control.

### 2.3. Microbial network modulation following EPF treatments

All treatments exhibited distinct microbial co-occurrence patterns (Fig. 5). The Mb network had the most nodes (70) but lowest density (0.0414), indicating high diversity with lower connectivity. Velifer had fewer nodes (52) but a higher density (0.0754), suggesting a more tightly connected structure, while the control was intermediate (63 nodes; 0.0754 density). Average path length and diameter reflected potential resource transfer: Mb (2.47; 7) was slowest, Velifer (1.70; 4) fastest, and the control intermediate (1.85; 5). Modularity was highest in Mb (0.795) compared to the control (0.598) and Velifer (0.585), suggesting stronger subcommunity structure. Dominant phyla differed: in control —*Bacteroidetes* (31.7%) and *Actinobacteria* (17.5%); in Mb — *Proteobacteria* (34.3%) and *Bacteroidetes* (18.6%), and in Velifer — *Bacteroidetes* (38.5%) and *Thaumarchaeota* (15.4%). All correlations were positive, indicating cooperative interactions. Mb showed the most significant correlations (9758) with Velifer showing fewer (5883). These results suggest that Mb promotes a diverse, modular community with strong positive associations, while Velifer fosters a smaller, more densely connected network with faster potential responses; the control remained intermediate **(Supplementary Tables 12–14**).

**Fig. 5.**
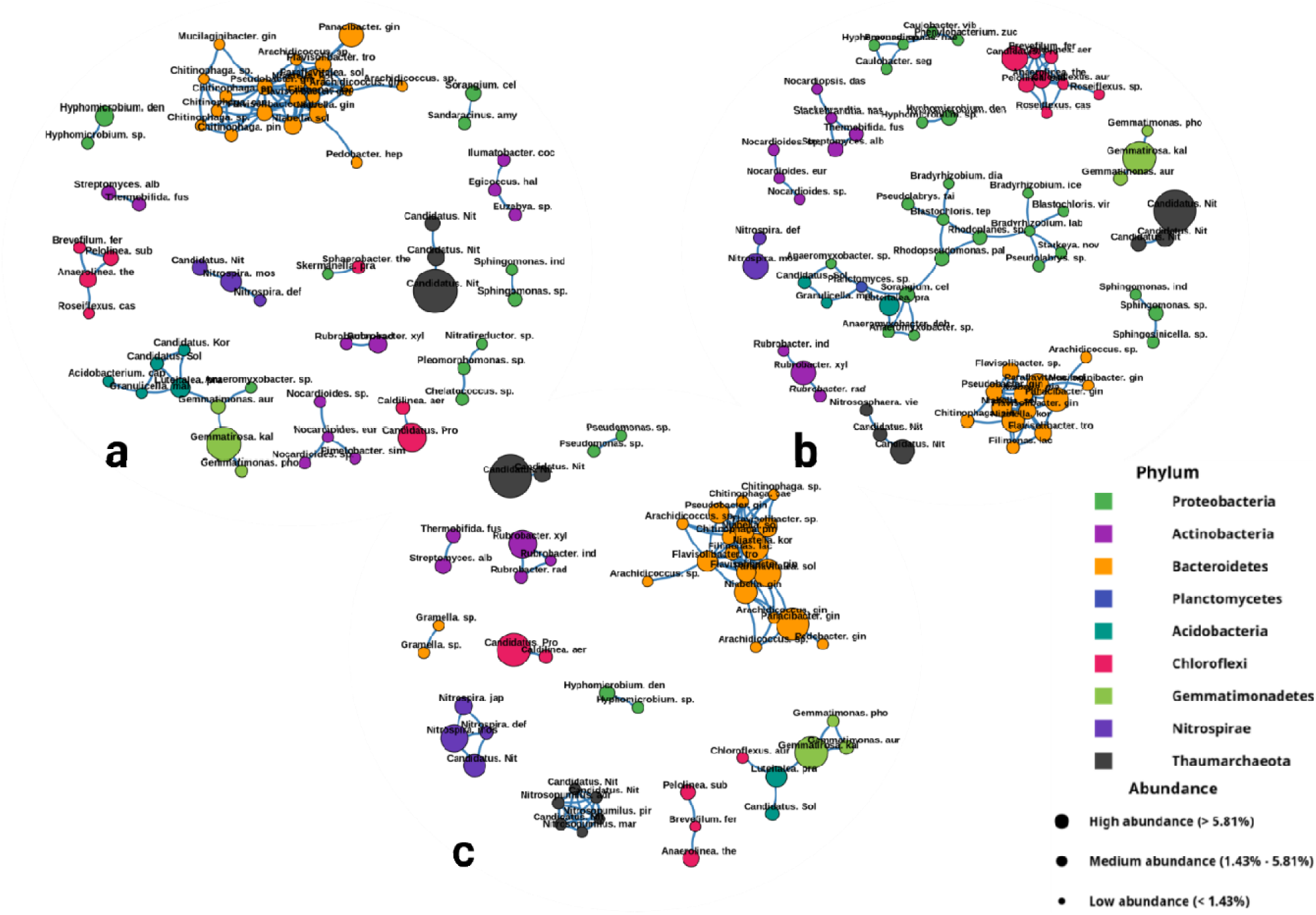
Microbial co-occurrence network analysis. The microbial co-occurrence networks for microbial communities under different treatments **a** Mb **b** control, and **c** Velifer; analyzed by SparCC analysis (R > 0. and *p* < 0.05); nodes represent species, colors represent phyla.

### 2.4. Functional Impacts of Mb and Velifer Treatments

Log2 fold-change analysis of microbial abundance between pre-application (−1) and 90 days post-application revealed treatment-specific impacts on soil microbial groups. Velifer was strongly associated with the nitrogen-fixing bacteria *Azotobacter* (logFC = 3.53), plant-beneficial *Pseudomonas* (logFC = 2.77), organic matter decomposers (mean logFC = 1.38 across 14 species), and antibiotic producers (*Amycolatopsis*, *Streptomyces* logFC = 1.88). Mb was associated with organic matter-decomposing bacteria such as *Anseongella ginsenosidimutans*, (logFC = 3.96), more moderately with *Pseudomonas* (logFC = 1.65), nitrogen-fixers (logFC = 1.06), denitrifiers (logFC = 1.05), and EPF abundance (logFC = 2.75). Overall, Velifer induced broader associations, whereas Mb caused narrower but stronger effects, including reduced denitrifier abundance (Fig. 6).

**Fig. 6.**
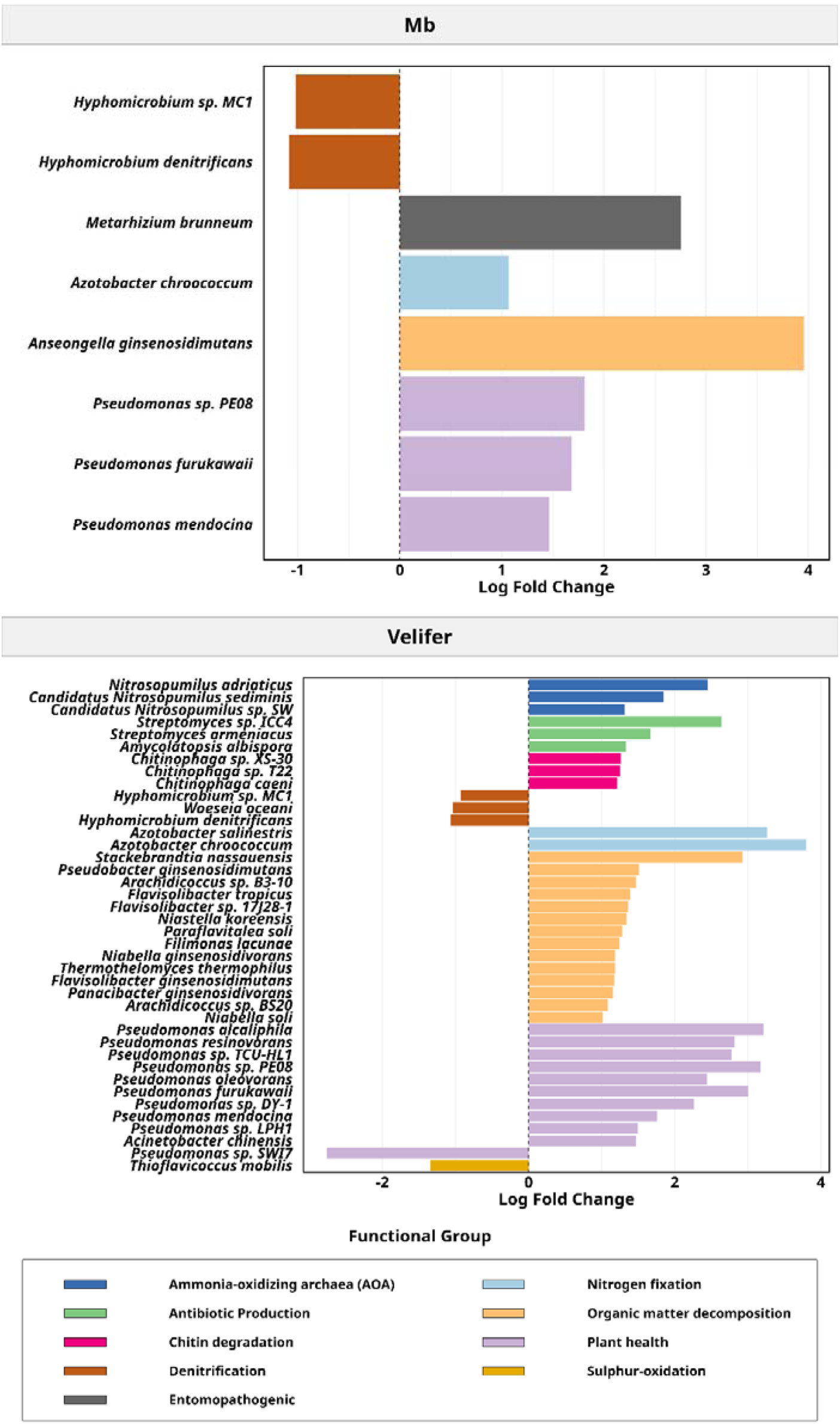
Log2 fold change analysis of differential species abundance with a significance threshold of *p* < 0.05 and functional annotation. Bars represent the magnitude and direction of abundance changes (log2 fold change), with positive values indicating increased abundance and negative values indicating decreased abundance, according to metabolic function.

## 3. Discussion

The trends in treatment efficacy suggest that Mb provides strong initial protection, whereas Velifer’s protective effect increases over time, suggesting potential for sustained efficacy. Mb and Velifer treatments were equally effective, with the most pronounced effect observed in soil, and maximal effect achieved immediately post-application remaining high at 180 days, confirming the stable and effective presence of EPF in soil.

Following the inundative application of EPF as a prophylactic myco-biopesticide in a commercial date plantation, we profiled palm-associated soil microbiota at four time points over a 90-day period. Our field design allowed us to link EPF to three main outcomes: palm health and protection against RPW infestation, entomopathogenic fungal persistence, and shifts in surrounding microbiota. Using the Galleria baiting method (GBM) methodology and metagenome analysis, we monitored the temporal persistence of EPF and native microbiota. Despite the limited number of sampled palms, the palm health index could be associated with microbial composition across repeated samplings, revealing that EPF persistence and specific community changes coincided with improved palm condition.

### 3.1. Palm protection and the role of EPF

Palm tree health was monitored over a 90-day period post-application using seismic sensors as reported before (Fig 1.c). The non-treated control palms exhibited fluctuations in health index, with a decline in palm health rates during the summer, aligning with the increased activity of the RPW as temperatures rise^15^. Velifer treatment provided consistent palm protection with minimal decline over time, whereas Mb treatment initially provided better protective effect (January–April), but infestation rates rose during the summer^15^. In terms of EPF efficacy, these findings suggest that Mb offers strong initial protection followed by a significant decline, whereas Velifer is more suitable for long-term application. Unlike the commercial product Velifer, Mb is a laboratory preparation of *M. brunneum* with minimal additives, which may have reduced protective properties against abiotic stress compared to commercial formulations ^16,17^. Therefore, a commercial formulation of *M. brunneum* can provide suitable protection against RPW as observed for other insects in laboratory and field conditions ^18,19^. Moreover, our previous results showed higher virulence of Mb toward RPW eggs compared to *B. bassiana*-based products in laboratory experiments^16,20^. However, the mechanism cannot be resolved under field conditions. Despite the presence of multiple native *Metarhizium* and *B. bassiana* in control soils, background EPF communities did not prevent RPW damage, suggesting that accessibility to the host, virulence, and inoculum density are critical for effective field protection^21,22^.

### 3.2. Factors affecting the background array of EPF

The composition of soil microbiota varied across samples, reflecting both treatment and site heterogeneity. Native *Metarhizium* spp. and *B. bassiana* confirmed in soils surrounding the palms and, as expected, demonstrated stable background fungal infectivity. Mb-treated plots showed late picks of *M. anisopliae* and *M. brunneum* but not *B. bassiana*, consistent with niche competition and the high stress tolerance reported for *Metarhizium* spp. ^23–25^. In Velifer plots, the late increase of *B. bassiana* suggests that niche conditions rather than inoculation alone governed proliferation. In contrast, control plots maintained a relatively even ratio between *M. anisopliae*, *M. brunneum*, and *B. bassiana* over time.

### 3.3. Geographical and temporal influence on the microbiota composition

Our results indicate that geography strongly influenced microbiota structure and potentially treatment response. Averaging triplicates confirmed strong separation by geography, suggesting EPF application had limited long-term effects. Shannon diversity supported these findings as it remained consistent across treatments. Yet, significant differences in Velifer indicated that *B. bassiana* application modulates soil microbial community, consistent with previous reports^6,26^. Simpson diversity and palm health data suggested dominance, time-dependent and seasonal effects, with EPF treatments showing great variability and reductions in diversity, a previously observed outcome in *M. brunneum*^17^. These treatment effects likely reflect a combination of competitive interactions and niche occupation by EPF and associated taxa, together with nutrient mediated shifts arising from changes in organic substrates and insect derived resources^2,6,27^.

### 3.4. Key interactions and species following EPF application

Bacterial relative abundance was stable across groups before application, but later shifts in major phyla, particularly *Proteobacteria*, *Bacteroidetes*, and *Actinobacteria* were observed. Mb increased plant-beneficial and nitrogen-fixing bacteria while reducing denitrifiers, suggesting a selective promotion of plant-growth and decomposer taxa that, to our knowledge, has not previously been reported at this high taxonomic resolution for field-applied EPF ^27–32^, Velifer, in contrast, drove broader functional shifts, plant-beneficial taxa, chitin degraders and antibiotic producers were enriched. This proliferation may be linked to the decline of *B. bassiana* relative abundance with time, which provides chitin as a substrate accompanied by chitin-rich insects digestion by the fungi ^33^. An interesting phenomenon was observed by the increase of nitrogen-fixers archaea under Velifer treatment, despite the known increase in nitrogen obtained by *Beauveria* insect-degradation^34^.

Mb exerted an immediate, selective influence—likely mediated by destruxins— favoring plant-growth promoters^35,36^. Velifer induced broader restructuring, enhancing functions across nutrient cycling, antibiotic production, and decomposition. Archaeal responses were diverged and support functional stability and adaptability, reinforcing Velifer’s potential in IPM. Beneficial taxa included diazotrophs, which fix N to NH via *nitrogenase*^37,38^ are crucial for soil fertility (ref). Co-inoculation of *B. bassiana* with nitrogen fixers is emerging as a sustainable biofertilizer strategy (ref). Ammonia-oxidizing archaea, which oxidize NH to NO, often outperform AOB in low-ammonia niches^39,40^, while denitrification remains largely bacterial (ref).

The significant decrease of pathogenic organism in treated soils demonstrates their role in plant protection^41^.

*B. bassiana* and *M. brunneum* not only support IPM but also enhance ecosystem functions (ref). The application of EPF to soil microbiota revealed that Velifer induces significant shifts in the abundance of a diverse range of species. Based on the variety of significantly changed species, the disturbance to the existing microbial community by Velifer application could be due to several mechanisms: enhancement of EPF that is relatively low abundance in an ecosystem could cause shifts in nutrient availability due to the decomposition of insect cadavers, triggering differences in the production of root exudates because *B. bassiana* is a rhizosphere colonizer^44^; this might contribute to higher availability of carbon sources and amino acids, which can be used advantageously by heterotrophic bacteria.

## 4. Methods

### 4.1. Experimental setup

The experiment was conducted in 2019, at Eden Farm Regional R&D in Bet-She’an Valley, Israel (32°28’01.8”N, 35°29’33.9”E; ITM X-246545.20730951097Y-708160.155770451). All of the palm trees were equipped with transmitting seismic sensors (Agrint Inc.), which constantly indicated the health index status for at least 1 month prior to the initiation of the experiment. Only healthy palms were chosen for the experiment (seismic-sensor-derived information from an area known to be infested with RPW). The experimental design consisted of 5 untreated trees (control group), 5 trees treated with the commercial product Velifer® (*Beauveria bassiana* PPRI 5339), and 5 trees treated with *Metarhizium brunneum* strain 7 (Mb), which was mass-produced in the laboratory as described previously ^17^. Fungal material was applied by spraying within the RPW cage trap surrounding the palm tree, 50 cm radius. The study area consisted of a rectangular plot, measuring 23.3 m in width and 170.6 m in length. The plot was further subdivided into sections to accommodate the different treatments (Fig. 1b).

### 4.2. Sample collection

Soil samples were collected from multiple sites around each tree before fungal application (day −1), at fungal application (day 0), and 30 and 90 days post-application. Samples were collected using sterile sampling tools at a depth of approximately 10 cm, and an approximate 50 cm radius around each tree, in triplicates. Care was taken to avoid surface debris.

### 4.3. Efficacy bioassay and single-spore isolation

Fungal persistence was tested using the *Galleria mellonella* bait (GMB) assay^45^. Soil samples were placed in 90-mm Petri dishes with 5 fifth-instar larvae and incubated at 25 °C until mortality. Infected larvae were surface-sterilized, transferred to filter paper in sterile Petri dishes, and incubated for sporulation. Spores were collected and single-spore isolates obtained by repeated transfers on SDA plates with chloramphenicol (10% v/w) and Dodine (16.25% v/w). Plates were incubated at 25 °C (*B. bassiana*) or 28 °C (*M. brunneum*), and multiple isolates were recovered to capture fungal diversity.

### 4.4. DNA extraction

Total DNA was extracted from soil using the DNeasy® PowerSoil® Kit (Qiagen) and quantified with a NanoDrop 2000. Fungi from larval cadavers were scraped from SDA plates and processed in a Spex Geno Grinder® with CTAB, β-mercaptoethanol, and RNase A^46^. Samples underwent incubation, chloroform:isoamyl alcohol extraction, DNA precipitation with Na acetate and isopropanol, ethanol washing, and SpeedVac™ drying. DNA was resuspended in ultrapure water, stored at −20 °C, and quantified/assessed for quality and contaminants by NanoDrop for Sanger sequencing.

### 4.5. Bioinformatic MetaWRAP pipeline

DNA was sequenced with Illumina NovaSeq 6000 (150-bp paired-end reads). Raw reads were processed with MetaWRAP (QC, assembly, binning, taxonomy, abundance)^47^. QC was done with FastQC; Trimmomatic trimmed low-quality bases (Phred ≥30, length ≥100 bp). Contaminants were filtered by mapping to RefSeq with Bowtie2, retaining unmapped reads (Bioproject PRJNA1273514). High-quality reads were assembled with MEGAHIT into 142,542 contigs; contigs ≥2000 bp (n=110,021) were binned with CONCOCT and refined with CheckM (≥50% completeness, ≤10% contamination), yielding 68 bins. Taxonomy was assigned with Kraken2. Abundance was normalized with DESeq2, and differential testing performed with edgeR (log-fold change, FDR q<0.05).

### 4.6. PCR amplification of the internal transcribed spacer (ITS) and elongation factor 1-alpha (EFT-1α) regions for phylogeny

The ITS region was amplified with general primers^48^ (Table 2) using PCRBIO HS Taq Mix Red. Cycling: 95 °C for 15 s, 35 cycles at 51 °C, final extension at 72 °C. The EF1-α gene was amplified under similar conditions with annealing at 53 °C^52^. PCR products were verified on 1% agarose gels, purified with the Zymo Gel DNA Recovery Kit, quantified on a NanoDrop 2000, and Sanger sequenced at Macrogen Europe. Chromatograms were analyzed in MEGA11^53^ and identified via NCBI BLAST.

**Table 2.**
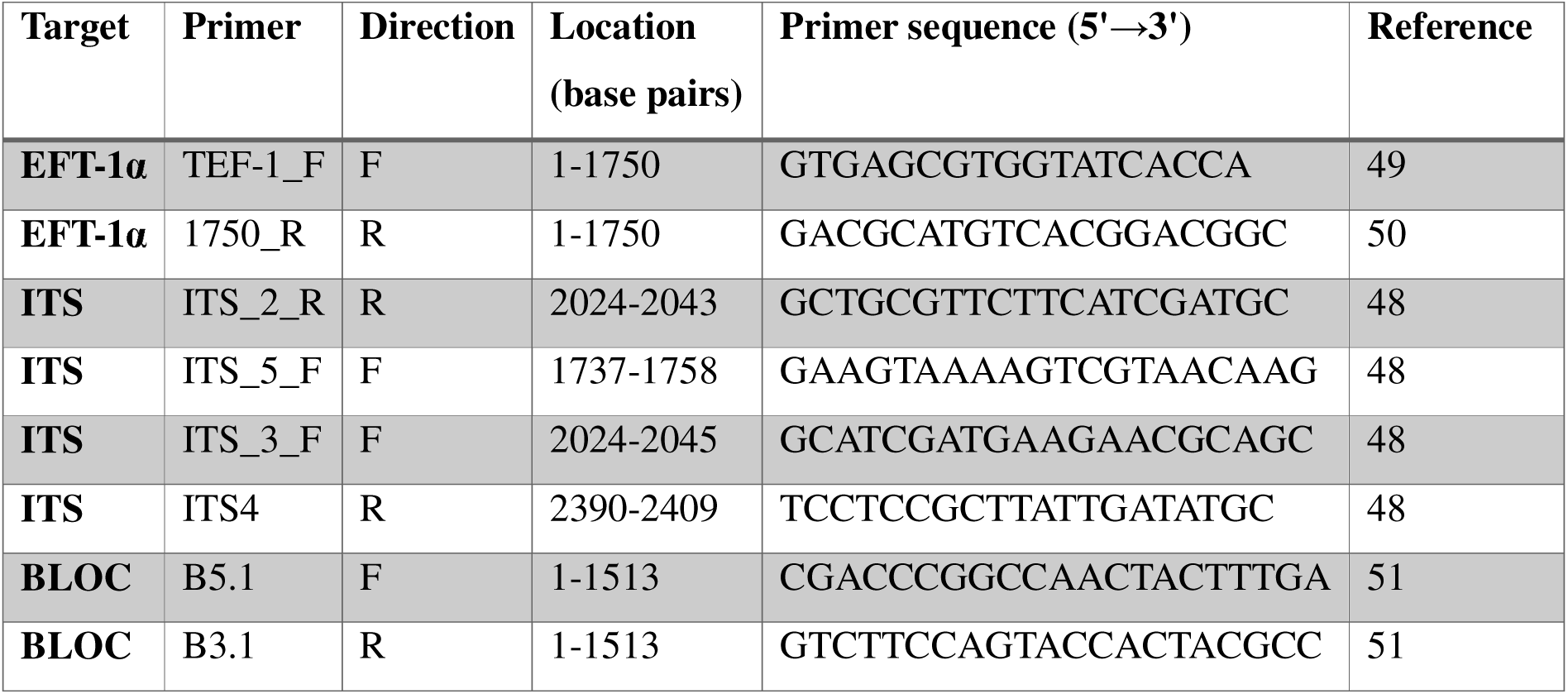
Primers used in this study.

### 4.7. Data analysis

For data analysis, contig abundances were aggregated at the species level. Species with counts <4 or prevalence <20% were removed to exclude rare/low-abundance noise. An interquartile variance filter excluded the bottom 10% least variable species. Data were rarefied to 159,145 contig counts and normalized using trimmed mean of M-values (TMM) to correct for compositional bias.

#### Alpha diversity

The Shannon diversity index indicates species richness and evenness, incorporating the relative abundance of each species: 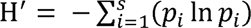. The Simpson’s diversity index considers both richness and evenness, emphasizing dominant species: 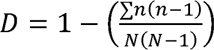.

#### Beta diversity

Temporal dynamics of microbial composition across treatments (control, Mb, Velifer) were assessed with NMDS and validated by PERMANOVA using the vegan R package (v2.6-4) on 42 samples. Data was pre-processed with Wisconsin double standardization, square-root transformation, and removal of zero-variance features. NMDS was performed in two dimensions with Bray–Curtis distance, 200 iterations, 100 random starts, and seed 123. PERMANOVA (999 unrestricted permutations, Bray–Curtis, treatment as main effect) tested group differences. Ordination was visualized in ggplot2 with 95% confidence ellipses per treatment.

#### Relative abundance and co-occurrence network

Relative abundance analysis of species was normalized with TMM via edgeR package and visualized with phyloseq, R Color Brewer packages in R. Abundance data were processed based on the SparCC algorithm^54^, with lowest correlation threshold set to 0.5 and significance threshold at *p* < 0.05.

## Supporting information

Matveev et al supplemental

## 5. Data availability

All data is available at Bioproject PRJNA1273514 and upon request from the corresponding author.

## 6. Acknowledgments

This study was funded by the Ministry of Agriculture and Rural Development and the Palm Growers Board (Grant no. 20-02-0091), and ICA in Israel (Grant no. 279/21) to DM. Our heartfelt appreciation goes to the Ment laboratory members for their continuous support and fruitful discussions.

## AUTHOR CONTRIBUTIONS

DM conceived and designed research. SM, VR, MD, YL conducted experiments. SM, VR, AF analyzed data. DM and SM wrote the manuscript. DM recruited funds. All authors read and approved the manuscript

## COMPETING INTERESTS

The authors have no competing interests.

## Notes

### Competing Interest Statement

The authors have declared no competing interest.

### Summary of Updates

We have revised the manuscript based on inputs we have received from a review process.

