## Supplementary material for "Field persistence of entomopathogenic fungi as biocontrol agents and network-level associations within the soil microbial community": Matveev et al supplemental

**Supplementary materials**

**Supplementary Table 1 | Tree sensor data indicating red palm weevil movement**

| **Tree** | **Treatment** | 7 Jan  2019 | 24 Jan 2019 | 6 Feb 2019 | 4 Mar 2019 | 10 Apr 2019 | 17 Apr 2019 | 1 May 2019 | 22 May 2019 | 12 Jun 2019 | 18 Jun 2019 | 9 Jul 2019 | 11 Aug 2019 |
| --- | --- | --- | --- | --- | --- | --- | --- | --- | --- | --- | --- | --- | --- |
| 41 | Mb | C | C | C | C | C | C | C | C | I | I | C | I |
| 42 | Control | C | C | C | C | C | C | C | C | I | C | C | I |
| 48 | Control | C | C | C | C | C | C | C | C | C | C | C | C |
| 49 | Control | C | C | C | C | C | C | C | C | C | C | C | C |
| 62 | Velifer | C | C | C | C | C | C | C | C | C | C | C | C |
| 63 | Mb | C | C | C | C | C | C | C | C | C | C | C | I |
| 67 | Control | C | C | C | C | C | C | C | C | C | C | C | C |
| 68 | Mb | C | C | C | C | C | C | C | C | C | C | C | C |
| 71 | Mb | C | C | C | C | C | C | C | C | I | C | C | C |
| 73 | Velifer | I | I | I | I | I | I | I | C | C | C | C | I |
| 75 | Velifer | C | C | C | C | C | C | C | C | C | C | C | C |
| 76 | Control | C | C | C | C | C | C | I | C | C | C | I | I |
| 77 | Velifer | C | C | C | C | I | I | C | C | C | C | C | C |
| 79 | Velifer | C | C | C | C | C | C | C | C | C | C | C | C |
| 80 | Mb | C | C | C | C | C | C | C | C | C | C | I | I |

Tree status: C, clean, no red palm weevil infestation; I, infested.

**Supplementary Table 2 | Full dataset for ordinary least squares** (**OLS) analysis**

| **Tree number** | **Treatment** | **Observation date**  **(dd/mm/yyyy)** | **Status** | **Health rate** | **Days** |
| --- | --- | --- | --- | --- | --- |
| 41 | Mb | 07/01/2019 | Clean | 100 | 0 |
| 41 | Mb | 24/01/2019 | Clean | 100 | 17 |
| 41 | Mb | 06/02/2019 | Clean | 100 | 30 |
| 41 | Mb | 04/03/2019 | Clean | 100 | 56 |
| 41 | Mb | 10/04/2019 | Clean | 100 | 93 |
| 41 | Mb | 17/04/2019 | Clean | 100 | 100 |
| 41 | Mb | 01/05/2019 | Clean | 100 | 114 |
| 41 | Mb | 22/05/2019 | Clean | 100 | 135 |
| 41 | Mb | 12/06/2019 | INFESTED | 0 | 156 |
| 41 | Mb | 18/06/2019 | INFESTED | 0 | 162 |
| 41 | Mb | 09/07/2019 | Clean | 100 | 183 |
| 41 | Mb | 11/08/2019 | INFESTED | 0 | 216 |
| 42 | Control | 07/01/2019 | Clean | 100 | 0 |
| 42 | Control | 24/01/2019 | Clean | 100 | 17 |
| 42 | Control | 06/02/2019 | Clean | 100 | 30 |
| 42 | Control | 04/03/2019 | Clean | 100 | 56 |
| 42 | Control | 10/04/2019 | Clean | 100 | 93 |
| 42 | Control | 17/04/2019 | Clean | 100 | 100 |
| 42 | Control | 01/05/2019 | Clean | 100 | 114 |
| 42 | Control | 22/05/2019 | Clean | 100 | 135 |
| 42 | Control | 12/06/2019 | INFESTED | 0 | 156 |
| 42 | Control | 18/06/2019 | Clean | 100 | 162 |
| 42 | Control | 09/07/2019 | Clean | 100 | 183 |
| 42 | Control | 11/08/2019 | INFESTED | 0 | 216 |
| 48 | Control | 07/01/2019 | Clean | 100 | 0 |
| 48 | Control | 24/01/2019 | Clean | 100 | 17 |
| 48 | Control | 06/02/2019 | Clean | 100 | 30 |
| 48 | Control | 04/03/2019 | Clean | 100 | 56 |
| 48 | Control | 10/04/2019 | Clean | 100 | 93 |
| 48 | Control | 17/04/2019 | Clean | 100 | 100 |
| 48 | Control | 01/05/2019 | Clean | 100 | 114 |
| 48 | Control | 22/05/2019 | Clean | 100 | 135 |
| 48 | Control | 12/06/2019 | Clean | 100 | 156 |
| 48 | Control | 18/06/2019 | Clean | 100 | 162 |
| 48 | Control | 09/07/2019 | Clean | 100 | 183 |
| 48 | Control | 11/08/2019 | Clean | 100 | 216 |
| 49 | Control | 07/01/2019 | Clean | 100 | 0 |
| 49 | Control | 24/01/2019 | Clean | 100 | 17 |
| 49 | Control | 06/02/2019 | Clean | 100 | 30 |
| 49 | Control | 04/03/2019 | Clean | 100 | 56 |
| 49 | Control | 10/04/2019 | Clean | 100 | 93 |
| 49 | Control | 17/04/2019 | Clean | 100 | 100 |
| 49 | Control | 01/05/2019 | Clean | 100 | 114 |
| 49 | Control | 22/05/2019 | Clean | 100 | 135 |
| 49 | Control | 12/06/2019 | Clean | 100 | 156 |
| 49 | Control | 18/06/2019 | Clean | 100 | 162 |
| 49 | Control | 09/07/2019 | Clean | 100 | 183 |
| 49 | Control | 11/08/2019 | Clean | 100 | 216 |
| 62 | Velifer | 07/01/2019 | Clean | 100 | 0 |
| 62 | Velifer | 24/01/2019 | Clean | 100 | 17 |
| 62 | Velifer | 06/02/2019 | Clean | 100 | 30 |
| 62 | Velifer | 04/03/2019 | Clean | 100 | 56 |
| 62 | Velifer | 10/04/2019 | Clean | 100 | 93 |
| 62 | Velifer | 17/04/2019 | Clean | 100 | 100 |
| 62 | Velifer | 01/05/2019 | Clean | 100 | 114 |
| 62 | Velifer | 22/05/2019 | Clean | 100 | 135 |
| 62 | Velifer | 12/06/2019 | Clean | 100 | 156 |
| 62 | Velifer | 18/06/2019 | Clean | 100 | 162 |
| 62 | Velifer | 09/07/2019 | Clean | 100 | 183 |
| 62 | Velifer | 11/08/2019 | Clean | 100 | 216 |
| 63 | Mb | 07/01/2019 | Clean | 100 | 0 |
| 63 | Mb | 24/01/2019 | Clean | 100 | 17 |
| 63 | Mb | 06/02/2019 | Clean | 100 | 30 |
| 63 | Mb | 04/03/2019 | Clean | 100 | 56 |
| 63 | Mb | 10/04/2019 | Clean | 100 | 93 |
| 63 | Mb | 17/04/2019 | Clean | 100 | 100 |
| 63 | Mb | 01/05/2019 | Clean | 100 | 114 |
| 63 | Mb | 22/05/2019 | Clean | 100 | 135 |
| 63 | Mb | 12/06/2019 | Clean | 100 | 156 |
| 63 | Mb | 18/06/2019 | Clean | 100 | 162 |
| 63 | Mb | 09/07/2019 | Clean | 100 | 183 |
| 63 | Mb | 11/08/2019 | INFESTED | 0 | 216 |
| 67 | Control | 07/01/2019 | Clean | 100 | 0 |
| 67 | Control | 24/01/2019 | Clean | 100 | 17 |
| 67 | Control | 06/02/2019 | Clean | 100 | 30 |
| 67 | Control | 04/03/2019 | Clean | 100 | 56 |
| 67 | Control | 10/04/2019 | Clean | 100 | 93 |
| 67 | Control | 17/04/2019 | Clean | 100 | 100 |
| 67 | Control | 01/05/2019 | Clean | 100 | 114 |
| 67 | Control | 22/05/2019 | Clean | 100 | 135 |
| 67 | Control | 12/06/2019 | Clean | 100 | 156 |
| 67 | Control | 18/06/2019 | Clean | 100 | 162 |
| 67 | Control | 09/07/2019 | Clean | 100 | 183 |
| 67 | Control | 11/08/2019 | Clean | 100 | 216 |
| 68 | Mb | 07/01/2019 | Clean | 100 | 0 |
| 68 | Mb | 24/01/2019 | Clean | 100 | 17 |
| 68 | Mb | 06/02/2019 | Clean | 100 | 30 |
| 68 | Mb | 04/03/2019 | Clean | 100 | 56 |
| 68 | Mb | 10/04/2019 | Clean | 100 | 93 |
| 68 | Mb | 17/04/2019 | Clean | 100 | 100 |
| 68 | Mb | 01/05/2019 | Clean | 100 | 114 |
| 68 | Mb | 22/05/2019 | Clean | 100 | 135 |
| 68 | Mb | 12/06/2019 | Clean | 100 | 156 |
| 68 | Mb | 18/06/2019 | Clean | 100 | 162 |
| 68 | Mb | 09/07/2019 | Clean | 100 | 183 |
| 68 | Mb | 11/08/2019 | Clean | 100 | 216 |
| 71 | Mb | 07/01/2019 | Clean | 100 | 0 |
| 71 | Mb | 24/01/2019 | Clean | 100 | 17 |
| 71 | Mb | 06/02/2019 | Clean | 100 | 30 |
| 71 | Mb | 04/03/2019 | Clean | 100 | 56 |
| 71 | Mb | 10/04/2019 | Clean | 100 | 93 |
| 71 | Mb | 17/04/2019 | Clean | 100 | 100 |
| 71 | Mb | 01/05/2019 | Clean | 100 | 114 |
| 71 | Mb | 22/05/2019 | Clean | 100 | 135 |
| 71 | Mb | 12/06/2019 | INFESTED | 0 | 156 |
| 71 | Mb | 18/06/2019 | Clean | 100 | 162 |
| 71 | Mb | 09/07/2019 | Clean | 100 | 183 |
| 71 | Mb | 11/08/2019 | Clean | 100 | 216 |
| 75 | Velifer | 07/01/2019 | Clean | 100 | 0 |
| 75 | Velifer | 24/01/2019 | Clean | 100 | 17 |
| 75 | Velifer | 06/02/2019 | Clean | 100 | 30 |
| 75 | Velifer | 04/03/2019 | Clean | 100 | 56 |
| 75 | Velifer | 10/04/2019 | Clean | 100 | 93 |
| 75 | Velifer | 17/04/2019 | Clean | 100 | 100 |
| 75 | Velifer | 01/05/2019 | Clean | 100 | 114 |
| 75 | Velifer | 22/05/2019 | Clean | 100 | 135 |
| 75 | Velifer | 12/06/2019 | Clean | 100 | 156 |
| 75 | Velifer | 18/06/2019 | Clean | 100 | 162 |
| 75 | Velifer | 09/07/2019 | Clean | 100 | 183 |
| 75 | Velifer | 11/08/2019 | Clean | 100 | 216 |
| 76 | Control | 07/01/2019 | Clean | 100 | 0 |
| 76 | Control | 24/01/2019 | Clean | 100 | 17 |
| 76 | Control | 06/02/2019 | Clean | 100 | 30 |
| 76 | Control | 04/03/2019 | Clean | 100 | 56 |
| 76 | Control | 10/04/2019 | Clean | 100 | 93 |
| 76 | Control | 17/04/2019 | Clean | 100 | 100 |
| 76 | Control | 01/05/2019 | INFESTED | 0 | 114 |
| 76 | Control | 22/05/2019 | Clean | 100 | 135 |
| 76 | Control | 12/06/2019 | Clean | 100 | 156 |
| 76 | Control | 18/06/2019 | Clean | 100 | 162 |
| 76 | Control | 09/07/2019 | INFESTED | 0 | 183 |
| 76 | Control | 11/08/2019 | INFESTED | 0 | 216 |
| 77 | Velifer | 07/01/2019 | Clean | 100 | 0 |
| 77 | Velifer | 24/01/2019 | Clean | 100 | 17 |
| 77 | Velifer | 06/02/2019 | Clean | 100 | 30 |
| 77 | Velifer | 04/03/2019 | Clean | 100 | 56 |
| 77 | Velifer | 10/04/2019 | INFESTED | 0 | 93 |
| 77 | Velifer | 17/04/2019 | INFESTED | 0 | 100 |
| 77 | Velifer | 01/05/2019 | Clean | 100 | 114 |
| 77 | Velifer | 22/05/2019 | Clean | 100 | 135 |
| 77 | Velifer | 12/06/2019 | Clean | 100 | 156 |
| 77 | Velifer | 18/06/2019 | Clean | 100 | 162 |
| 77 | Velifer | 09/07/2019 | Clean | 100 | 183 |
| 77 | Velifer | 11/08/2019 | Clean | 100 | 216 |
| 79 | Velifer | 07/01/2019 | Clean | 100 | 0 |
| 79 | Velifer | 24/01/2019 | Clean | 100 | 17 |
| 79 | Velifer | 06/02/2019 | Clean | 100 | 30 |
| 79 | Velifer | 04/03/2019 | Clean | 100 | 56 |
| 79 | Velifer | 10/04/2019 | Clean | 100 | 93 |
| 79 | Velifer | 17/04/2019 | Clean | 100 | 100 |
| 79 | Velifer | 01/05/2019 | Clean | 100 | 114 |
| 79 | Velifer | 22/05/2019 | Clean | 100 | 135 |
| 79 | Velifer | 12/06/2019 | Clean | 100 | 156 |
| 79 | Velifer | 18/06/2019 | Clean | 100 | 162 |
| 79 | Velifer | 09/07/2019 | Clean | 100 | 183 |
| 79 | Velifer | 11/08/2019 | Clean | 100 | 216 |
| 80 | Mb | 07/01/2019 | Clean | 100 | 0 |
| 80 | Mb | 24/01/2019 | Clean | 100 | 17 |
| 80 | Mb | 06/02/2019 | Clean | 100 | 30 |
| 80 | Mb | 04/03/2019 | Clean | 100 | 56 |
| 80 | Mb | 10/04/2019 | Clean | 100 | 93 |
| 80 | Mb | 17/04/2019 | Clean | 100 | 100 |
| 80 | Mb | 01/05/2019 | Clean | 100 | 114 |
| 80 | Mb | 22/05/2019 | Clean | 100 | 135 |
| 80 | Mb | 12/06/2019 | Clean | 100 | 156 |
| 80 | Mb | 18/06/2019 | Clean | 100 | 162 |
| 80 | Mb | 09/07/2019 | INFESTED | 0 | 183 |
| 80 | Mb | 11/08/2019 | INFESTED | 0 | 216 |

**Supplementary Table 3 | Health rate by treatment date**

| **Observation date**  **(dd/mm/yyyy)** | **Control** | **Mb** | **Velifer** |
| --- | --- | --- | --- |
| 07/01/2019 | 100 | 100 | 100 |
| 24/01/2019 | 100 | 100 | 100 |
| 06/02/2019 | 100 | 100 | 100 |
| 04/03/2019 | 100 | 100 | 100 |
| 10/04/2019 | 100 | 100 | 75 |
| 17/04/2019 | 100 | 100 | 75 |
| 01/05/2019 | 80 | 100 | 100 |
| 22/05/2019 | 100 | 100 | 100 |
| 12/06/2019 | 80 | 60 | 100 |
| 18/06/2019 | 100 | 80 | 100 |
| 09/07/2019 | 80 | 80 | 100 |
| 11/08/2019 | 60 | 40 | 100 |

**Supplementary Table 4 | OLS model results**

| **Variable** | **Coefficient** | **SE** | **t_value** | ***p*_value** | **CI_lower** | **CI_upper** |
| --- | --- | --- | --- | --- | --- | --- |
| Intercept | 78.08828201 | 2.846898 | 27.4292546 | 9.09E-63 | 72.46647 | 83.7101 |
| Treatment_Control | 28.05064608 | 5.296713 | 5.295859143 | 3.81E-07 | 17.59114 | 38.51015 |
| Treatment_Mb | 33.16562127 | 5.296713 | 6.261547707 | 3.29E-09 | 22.70612 | 43.62512 |
| Treatment_Velifer | 16.87201467 | 5.748283 | 2.935140015 | 0.00381879 | 5.52079 | 28.22324 |
| Days | -0.086814092 | 0.022934 | -3.785468916 | 0.000215709 | -0.1321 | -0.04153 |
| Treatment_Control_Days | -0.050798536 | 0.042668 | -1.190545713 | 0.235573592 | -0.13506 | 0.033459 |
| Treatment_Mb_Days | -0.131131106 | 0.042668 | -3.073269167 | 0.002484803 | -0.21539 | -0.04687 |
| Treatment_Velifer_Days | 0.09511555 | 0.046306 | 2.054067548 | 0.041576098 | 0.003674 | 0.186557 |

**Supplementary Table 5 | Fungal treatment efficacy data**

| **Days relative to application (day 0)** | **Samp-ling season** | **Control GBM, % mortality, mean** | **Control GBM, % mortality,**  **SE** | **Control CFU/g dry, mean** | **Control CFU/g dry, SE** | **Mb GBM, % mortality, mean** | **Mb GBM, % mortality,**  **SE** | **Mb CFU/g dry, mean** | **Mb CFU/g dry, SE** | **Velifer GBM, % mortality, mean** | **Velifer GBM, % mortality,**  **SE** | **Velifer CFU/g dry,**  **mean** | **Velifer CFU/g dry,**  **SE** |
| --- | --- | --- | --- | --- | --- | --- | --- | --- | --- | --- | --- | --- | --- |
| -1 | Winter | 28.8 | 0.08 | 326.63 | 147 | 1.6 | 0.01 | 156.54 | 108.46 | 0 | 0 | 37.83 | 37.83 |
| 0 | Winter | 54.4 | 0.09 | 85.79 | 49.75 | 54.4 | 0.09 | 85.79 | 49.75 | 0.68 | 0.08 | 52750 | 6264.52 |
| 30 | Winter | 22.4 | 0.06 | 1766.6 | 694.45 | 44 | 0.07 | 6681.9 | 6713.5 | 0.5 | 0.08 | 78622.19 | 18071.8 |
| 90 | Spring | 18.4 | 0.06 | 626.94 | 264.8 | 63.2 | 0.05 | 1082.6 | 449.28 |  |  |  |  |
| 180 | Summer | 32.8 | 0.07 | 1799.45 | 802.65 | 69.6 | 0.08 | 16620 | 7148.9 | 0.58 | 0.09 | 41688.75 | 8112.2 |

GBM, *Galleria mellonella* bait method; CFU/g dry, colony-forming units pergram dry weight.

**Supplementary Table 6 | ANOVA-determined statistical effects for *Galleria mellonella* bait method**

| **Source** | **N parameters estimated** | **DF** | **Sum of squares** | **F ratio** | **Prob F** |
| --- | --- | --- | --- | --- | --- |
| Time relative to day of application | 4 | 4 | 7.5376155 | 14.4831 | <.0001* |
| Treatment | 2 | 2 | 5.6774744 | 21.8179 | <.0001* |
| Sample type | 1 | 1 | 4.2007692 | 32.2861 | <.0001* |

Day relative to day of application (day 0),

Student’s t-test:

| **Level** |  |  |  | **LSM** |
| --- | --- | --- | --- | --- |
| 0 | A |  |  | 0.43553333 |
| 180 | A | B |  | 0.37866667 |
| 90 | A | B |  | 0.37466667 |
| 30 |  | B |  | 0.30400000 |
| -1 |  |  | C | 0.14533333 |

Treatment, Student’s t-test:

| **Level** |  |  | **LSM** |
| --- | --- | --- | --- |
| Mb | A |  | 0.39520000 |
| Velifer | A |  | 0.38292000 |
| Control |  | B | 0.20480000 |

Sample type, Student’s t-test:

| **Level** |  |  | **LSM** |
| --- | --- | --- | --- |
| Soil | A |  | 0.40248000 |
| Tree bark |  | B | 0.25280000 |

LSM, least squares mean.

**Supplementary Table 7 | *Galleria mellonella* bait method assay results over time**

| **Days relative to day of application (day 0)** | **Sampling season** | | | **Treatment** | | | | | |
| --- | --- | --- | --- | --- | --- | --- | --- | --- | --- |
|  |  |  |  | Control |  | Mb |  | Velifer |  |
|  | Winter | Spring | Summer | Mean | SE | Mean | SE | Mean | SE |
| -1 | 0 |  |  | 28.80% | 7.75% | 1.60% | 1.11% | 1.50% | 0.00% |
| 0 | 0 |  |  | 28.80% | 7.75% | 54.40% | 8.91% | 67.70% | 8.21% |
| 30 | 0 |  |  | 22.40% | 5.92% | 44.00% | 7.39% | 50.40% | 7.84% |
| 90 |  | 0 |  | 18.40% | 5.88% | 63.20% | 4.57% | . | . |
| 180 |  |  | 0 | 32.80% | 7.20% | 69.60% | 7.58% | 57.60% | 9.12% |

**Supplementary Table 8 | Bacterial relative abundance with time and treatment**

| **Taxon** | **Time** | **Treatment** | **avg_rel_abundance** |
| --- | --- | --- | --- |
| *Azotobacter chroococcum* | -1 | Control | 2.486778307 |
| *Azotobacter chroococcum* | -1 | Mb | 6.20560515 |
| *Azotobacter chroococcum* | -1 | Velifer | 3.095573992 |
| *Azotobacter chroococcum* | 0 | Mb | 5.161242654 |
| *Azotobacter chroococcum* | 0 | Velifer | 5.898486034 |
| *Azotobacter chroococcum* | 30 | Control | 7.620195167 |
| *Azotobacter chroococcum* | 30 | Mb | 9.201824461 |
| *Azotobacter chroococcum* | 30 | Velifer | 8.296131801 |
| *Azotobacter chroococcum* | 90 | Control | 3.010786357 |
| *Azotobacter chroococcum* | 90 | Mb | 5.133253358 |
| *Azotobacter chroococcum* | 90 | Velifer | 8.732743078 |
| *Candidatus Promineofilum breve* | -1 | Control | 13.11501207 |
| *Candidatus Promineofilum breve* | -1 | Mb | 11.19597989 |
| *Candidatus Promineofilum breve* | -1 | Velifer | 14.25774501 |
| *Candidatus Promineofilum breve* | 0 | Mb | 10.42554167 |
| *Candidatus Promineofilum breve* | 0 | Velifer | 10.57132583 |
| *Candidatus Promineofilum breve* | 30 | Control | 12.36298704 |
| *Candidatus Promineofilum breve* | 30 | Mb | 8.273244288 |
| *Candidatus Promineofilum breve* | 30 | Velifer | 7.546838814 |
| *Candidatus Promineofilum breve* | 90 | Control | 15.32285859 |
| *Candidatus Promineofilum breve* | 90 | Mb | 10.45545926 |
| *Candidatus Promineofilum breve* | 90 | Velifer | 14.50706979 |
| *Chryseolinea soli* | -1 | Control | 10.75180915 |
| *Chryseolinea soli* | -1 | Mb | 10.45350161 |
| *Chryseolinea soli* | -1 | Velifer | 16.58030041 |
| *Chryseolinea soli* | 0 | Mb | 13.92594598 |
| *Chryseolinea soli* | 0 | Velifer | 15.34106191 |
| *Chryseolinea soli* | 30 | Control | 16.21177401 |
| *Chryseolinea soli* | 30 | Mb | 15.2559521 |
| *Chryseolinea soli* | 30 | Velifer | 10.97822099 |
| *Chryseolinea soli* | 90 | Control | 20.92990094 |
| *Chryseolinea soli* | 90 | Mb | 16.10039031 |
| *Chryseolinea soli* | 90 | Velifer | 14.99515047 |
| *Gemmatirosa kalamazoonesis* | -1 | Control | 18.98984932 |
| *Gemmatirosa kalamazoonesis* | -1 | Mb | 15.24610283 |
| *Gemmatirosa kalamazoonesis* | -1 | Velifer | 12.53719142 |
| *Gemmatirosa kalamazoonesis* | 0 | Mb | 16.28929839 |
| *Gemmatirosa kalamazoonesis* | 0 | Velifer | 12.12094455 |
| *Gemmatirosa kalamazoonesis* | 30 | Control | 19.50995865 |
| *Gemmatirosa kalamazoonesis* | 30 | Mb | 13.73631914 |
| *Gemmatirosa kalamazoonesis* | 30 | Velifer | 13.0630682 |
| *Gemmatirosa kalamazoonesis* | 90 | Control | 21.74924625 |
| *Gemmatirosa kalamazoonesis* | 90 | Mb | 15.13403098 |
| *Gemmatirosa kalamazoonesis* | 90 | Velifer | 11.78275713 |
| *Niabella soli* | -1 | Control | 7.016555089 |
| *Niabella soli* | -1 | Mb | 7.948540654 |
| *Niabella soli* | -1 | Velifer | 6.768020556 |
| *Niabella soli* | 0 | Mb | 7.270185256 |
| *Niabella soli* | 0 | Velifer | 8.097797431 |
| *Niabella soli* | 30 | Control | 6.57190745 |
| *Niabella soli* | 30 | Mb | 7.184946897 |
| *Niabella soli* | 30 | Velifer | 7.995309365 |
| *Niabella soli* | 90 | Control | 5.52822808 |
| *Niabella soli* | 90 | Mb | 5.42573567 |
| *Niabella soli* | 90 | Velifer | 6.561756306 |
| *Niastella koreensis* | -1 | Control | 8.489613941 |
| *Niastella koreensis* | -1 | Mb | 8.859498082 |
| *Niastella koreensis* | -1 | Velifer | 7.612096332 |
| *Niastella koreensis* | 0 | Mb | 8.188347857 |
| *Niastella koreensis* | 0 | Velifer | 9.410630404 |
| *Niastella koreensis* | 30 | Control | 7.921116566 |
| *Niastella koreensis* | 30 | Mb | 9.185497132 |
| *Niastella koreensis* | 30 | Velifer | 10.65589985 |
| *Niastella koreensis* | 90 | Control | 5.148785312 |
| *Niastella koreensis* | 90 | Mb | 7.056290006 |
| *Niastella koreensis* | 90 | Velifer | 8.134634338 |
| *Nitrospira moscoviensis* | -1 | Control | 11.40977633 |
| *Nitrospira moscoviensis* | -1 | Mb | 13.41492298 |
| *Nitrospira moscoviensis* | -1 | Velifer | 13.33038205 |
| *Nitrospira moscoviensis* | 0 | Mb | 14.28068211 |
| *Nitrospira moscoviensis* | 0 | Velifer | 11.48356919 |
| *Nitrospira moscoviensis* | 30 | Control | 6.489280434 |
| *Nitrospira moscoviensis* | 30 | Mb | 6.86715659 |
| *Nitrospira moscoviensis* | 30 | Velifer | 7.978605601 |
| *Nitrospira moscoviensis* | 90 | Control | 8.317813088 |
| *Nitrospira moscoviensis* | 90 | Mb | 8.283497041 |
| *Nitrospira moscoviensis* | 90 | Velifer | 6.491974075 |
| *Panacibacter ginsenosidivorans* | -1 | Control | 13.80589384 |
| *Panacibacter ginsenosidivorans* | -1 | Mb | 10.00026605 |
| *Panacibacter ginsenosidivorans* | -1 | Velifer | 10.11277322 |
| *Panacibacter ginsenosidivorans* | 0 | Mb | 10.11962226 |
| *Panacibacter ginsenosidivorans* | 0 | Velifer | 11.32883242 |
| *Panacibacter ginsenosidivorans* | 30 | Control | 10.30942543 |
| *Panacibacter ginsenosidivorans* | 30 | Mb | 11.9566465 |
| *Panacibacter ginsenosidivorans* | 30 | Velifer | 13.85800593 |
| *Panacibacter ginsenosidivorans* | 90 | Control | 6.237193364 |
| *Panacibacter ginsenosidivorans* | 90 | Mb | 8.41950028 |
| *Panacibacter ginsenosidivorans* | 90 | Velifer | 9.121652674 |
| *Paraflavitalea soli* | -1 | Control | 9.740191613 |
| *Paraflavitalea soli* | -1 | Mb | 9.396626064 |
| *Paraflavitalea soli* | -1 | Velifer | 8.630192752 |
| *Paraflavitalea soli* | 0 | Mb | 8.282547215 |
| *Paraflavitalea soli* | 0 | Velifer | 9.806590649 |
| *Paraflavitalea soli* | 30 | Control | 8.209153976 |
| *Paraflavitalea soli* | 30 | Mb | 9.327170832 |
| *Paraflavitalea soli* | 30 | Velifer | 10.64568829 |
| *Paraflavitalea soli* | 90 | Control | 5.15284387 |
| *Paraflavitalea soli* | 90 | Mb | 6.470874009 |
| *Paraflavitalea soli* | 90 | Velifer | 8.244730424 |
| *Rubrobacter xylanophilus* | -1 | Control | 4.194520344 |
| *Rubrobacter xylanophilus* | -1 | Mb | 7.278956684 |
| *Rubrobacter xylanophilus* | -1 | Velifer | 7.075724261 |
| *Rubrobacter xylanophilus* | 0 | Mb | 6.056586612 |
| *Rubrobacter xylanophilus* | 0 | Velifer | 5.940761575 |
| *Rubrobacter xylanophilus* | 30 | Control | 4.794201277 |
| *Rubrobacter xylanophilus* | 30 | Mb | 9.011242061 |
| *Rubrobacter xylanophilus* | 30 | Velifer | 8.982231166 |
| *Rubrobacter xylanophilus* | 90 | Control | 8.602344147 |
| *Rubrobacter xylanophilus* | 90 | Mb | 17.52096909 |
| *Rubrobacter xylanophilus* | 90 | Velifer | 11.42753172 |

Time: -1, 1 day before treatment application; 0, day of treatment application; 30 and 90, 30 and 90 days post-application.

**Supplementary Table 9 | Fungal relative abundance with time and treatment**

| **Taxon** | **Time** | **Treatment** | **avg_rel_abundance** |
| --- | --- | --- | --- |
| *Alternaria solani* | -1 | Control | 7.784498994 |
| *Alternaria solani* | -1 | Mb | 8.905223844 |
| *Alternaria solani* | -1 | Velifer | 7.901853822 |
| *Alternaria solani* | 0 | Mb | 6.844066851 |
| *Alternaria solani* | 0 | Velifer | 10.96548144 |
| *Alternaria solani* | 30 | Control | 6.24683707 |
| *Alternaria solani* | 30 | Mb | 11.60551023 |
| *Alternaria solani* | 30 | Velifer | 10.51637167 |
| *Alternaria solani* | 90 | Control | 8.233438875 |
| *Alternaria solani* | 90 | Mb | 7.980416752 |
| *Alternaria solani* | 90 | Velifer | 5.855615447 |
| *Fusarium venenatum* | -1 | Control | 1.724947906 |
| *Fusarium venenatum* | -1 | Mb | 1.361494706 |
| *Fusarium venenatum* | -1 | Velifer | 1.572515082 |
| *Fusarium venenatum* | 0 | Mb | 3.088058064 |
| *Fusarium venenatum* | 0 | Velifer | 2.765896313 |
| *Fusarium venenatum* | 30 | Control | 3.29839448 |
| *Fusarium venenatum* | 30 | Mb | 3.858963927 |
| *Fusarium venenatum* | 30 | Velifer | 3.344013458 |
| *Fusarium venenatum* | 90 | Control | 5.004204548 |
| *Fusarium venenatum* | 90 | Mb | 5.680812629 |
| *Fusarium venenatum* | 90 | Velifer | 6.257990705 |
| *Metarhizium anisopliae* | -1 | Control | 1.60485179 |
| *Metarhizium anisopliae* | -1 | Mb | 1.73526642 |
| *Metarhizium anisopliae* | -1 | Velifer | 1.042625726 |
| *Metarhizium anisopliae* | 0 | Mb | 7.230228251 |
| *Metarhizium anisopliae* | 0 | Velifer | 0.914819208 |
| *Metarhizium anisopliae* | 30 | Control | 2.321942333 |
| *Metarhizium anisopliae* | 30 | Mb | 6.111342149 |
| *Metarhizium anisopliae* | 30 | Velifer | 1.960372423 |
| *Metarhizium anisopliae* | 90 | Control | 3.317878483 |
| *Metarhizium anisopliae* | 90 | Mb | 6.040535713 |
| *Metarhizium anisopliae* | 90 | Velifer | 2.813959664 |
| *Metarhizium brunneum* | -1 | Control | 4.545731007 |
| *Metarhizium brunneum* | -1 | Mb | 4.22750123 |
| *Metarhizium brunneum* | -1 | Velifer | 4.209738776 |
| *Metarhizium brunneum* | 0 | Mb | 18.5575175 |
| *Metarhizium brunneum* | 0 | Velifer | 4.419103857 |
| *Metarhizium brunneum* | 30 | Control | 2.396452516 |
| *Metarhizium brunneum* | 30 | Mb | 11.74482141 |
| *Metarhizium brunneum* | 30 | Velifer | 3.428191936 |
| *Metarhizium brunneum* | 90 | Control | 2.529791083 |
| *Metarhizium brunneum* | 90 | Mb | 9.249374402 |
| *Metarhizium brunneum* | 90 | Velifer | 2.92938046 |
| *Mycosphaerella hyperici* | -1 | Control | 16.80480478 |
| *Mycosphaerella hyperici* | -1 | Mb | 17.93273324 |
| *Mycosphaerella hyperici* | -1 | Velifer | 23.13887453 |
| *Mycosphaerella hyperici* | 0 | Mb | 19.62423522 |
| *Mycosphaerella hyperici* | 0 | Velifer | 25.59721296 |
| *Mycosphaerella hyperici* | 30 | Control | 18.36055283 |
| *Mycosphaerella hyperici* | 30 | Mb | 20.36196658 |
| *Mycosphaerella hyperici* | 30 | Velifer | 24.91424418 |
| *Mycosphaerella hyperici* | 90 | Control | 22.9123315 |
| *Mycosphaerella hyperici* | 90 | Mb | 18.01010747 |
| *Mycosphaerella hyperici* | 90 | Velifer | 19.15164698 |
| *Parastagonospora nodorum* | -1 | Control | 4.097413639 |
| *Parastagonospora nodorum* | -1 | Mb | 4.626105508 |
| *Parastagonospora nodorum* | -1 | Velifer | 1.431001307 |
| *Parastagonospora nodorum* | 0 | Mb | 4.468617785 |
| *Parastagonospora nodorum* | 0 | Velifer | 4.710172533 |
| *Parastagonospora nodorum* | 30 | Control | 4.219794152 |
| *Parastagonospora nodorum* | 30 | Mb | 5.540453032 |
| *Parastagonospora nodorum* | 30 | Velifer | 4.195589181 |
| *Parastagonospora nodorum* | 90 | Control | 3.867933927 |
| *Parastagonospora nodorum* | 90 | Mb | 6.398911343 |
| *Parastagonospora nodorum* | 90 | Velifer | 1.999810447 |
| *Pertusaria propinqua* | -1 | Control | 14.16185778 |
| *Pertusaria propinqua* | -1 | Mb | 9.646434162 |
| *Pertusaria propinqua* | -1 | Velifer | 11.55304132 |
| *Pertusaria propinqua* | 0 | Mb | 4.231768199 |
| *Pertusaria propinqua* | 0 | Velifer | 8.529234931 |
| *Pertusaria propinqua* | 30 | Control | 7.428473597 |
| *Pertusaria propinqua* | 30 | Mb | 4.966729469 |
| *Pertusaria propinqua* | 30 | Velifer | 8.997927337 |
| *Pertusaria propinqua* | 90 | Control | 5.808843363 |
| *Pertusaria propinqua* | 90 | Mb | 3.2275325 |
| *Pertusaria propinqua* | 90 | Velifer | 4.877104237 |
| *Rhizopus oryzae* | -1 | Control | 5.674664431 |
| *Rhizopus oryzae* | -1 | Mb | 8.737746735 |
| *Rhizopus oryzae* | -1 | Velifer | 5.725312089 |
| *Rhizopus oryzae* | 0 | Mb | 4.809893828 |
| *Rhizopus oryzae* | 0 | Velifer | 7.960771632 |
| *Rhizopus oryzae* | 30 | Control | 6.904737554 |
| *Rhizopus oryzae* | 30 | Mb | 6.327138285 |
| *Rhizopus oryzae* | 30 | Velifer | 7.628090682 |
| *Rhizopus oryzae* | 90 | Control | 5.600548155 |
| *Rhizopus oryzae* | 90 | Mb | 5.968807081 |
| *Rhizopus oryzae* | 90 | Velifer | 4.933815977 |
| *Saprolegnia parasitica* | -1 | Control | 15.3602684 |
| *Saprolegnia parasitica* | -1 | Mb | 13.01476819 |
| *Saprolegnia parasitica* | -1 | Velifer | 4.874810027 |
| *Saprolegnia parasitica* | 0 | Mb | 4.788633168 |
| *Saprolegnia parasitica* | 0 | Velifer | 3.521884626 |
| *Saprolegnia parasitica* | 30 | Control | 5.558254534 |
| *Saprolegnia parasitica* | 30 | Mb | 3.102546116 |
| *Saprolegnia parasitica* | 30 | Velifer | 1.663514759 |
| *Saprolegnia parasitica* | 90 | Control | 5.512316937 |
| *Saprolegnia parasitica* | 90 | Mb | 3.602343874 |
| *Saprolegnia parasitica* | 90 | Velifer | 1.976336587 |
| *Thermothelomyces thermophilus* | -1 | Control | 28.24096128 |
| *Thermothelomyces thermophilus* | -1 | Mb | 29.81272596 |
| *Thermothelomyces thermophilus* | -1 | Velifer | 38.55022732 |
| *Thermothelomyces thermophilus* | 0 | Mb | 26.35698113 |
| *Thermothelomyces thermophilus* | 0 | Velifer | 30.6154225 |
| *Thermothelomyces thermophilus* | 30 | Control | 43.26456093 |
| *Thermothelomyces thermophilus* | 30 | Mb | 26.3805288 |
| *Thermothelomyces thermophilus* | 30 | Velifer | 33.35168438 |
| *Thermothelomyces thermophilus* | 90 | Control | 37.21271313 |
| *Thermothelomyces thermophilus* | 90 | Mb | 33.84115824 |
| *Thermothelomyces thermophilus* | 90 | Velifer | 49.2043395 |

Time: -1, 1 day before treatment application; 0, day of treatment application; 30 and 90, 30 and 90 days post-application.

**Supplementary Table 10 | Archaea relative abundance with time and treatment**

| **Taxon** | **Time** | **Treatment** | **avg_rel_abundance** |
| --- | --- | --- | --- |
| *Candidatus Nitrosocosmicus exaquare* | -1 | Control | 33.88936785 |
| *Candidatus Nitrosocosmicus exaquare* | -1 | Mb | 34.48803005 |
| *Candidatus Nitrosocosmicus exaquare* | -1 | Velifer | 29.03968628 |
| *Candidatus Nitrosocosmicus exaquare* | 0 | Mb | 40.30422226 |
| *Candidatus Nitrosocosmicus exaquare* | 0 | Velifer | 31.70787615 |
| *Candidatus Nitrosocosmicus exaquare* | 30 | Control | 45.41534285 |
| *Candidatus Nitrosocosmicus exaquare* | 30 | Mb | 36.13653073 |
| *Candidatus Nitrosocosmicus exaquare* | 30 | Velifer | 30.34110183 |
| *Candidatus Nitrosocosmicus exaquare* | 90 | Control | 59.80338961 |
| *Candidatus Nitrosocosmicus exaquare* | 90 | Mb | 49.78138464 |
| *Candidatus Nitrosocosmicus exaquare* | 90 | Velifer | 52.67018857 |
| *Candidatus Nitrosocosmicus franklandus* | -1 | Control | 9.069295438 |
| *Candidatus Nitrosocosmicus franklandus* | -1 | Mb | 9.803333401 |
| *Candidatus Nitrosocosmicus franklandus* | -1 | Velifer | 6.599910795 |
| *Candidatus Nitrosocosmicus franklandus* | 0 | Mb | 9.091618307 |
| *Candidatus Nitrosocosmicus franklandus* | 0 | Velifer | 6.907730444 |
| *Candidatus Nitrosocosmicus franklandus* | 30 | Control | 9.811127426 |
| *Candidatus Nitrosocosmicus franklandus* | 30 | Mb | 10.37481916 |
| *Candidatus Nitrosocosmicus franklandus* | 30 | Velifer | 7.483569305 |
| *Candidatus Nitrosocosmicus franklandus* | 90 | Control | 12.17566562 |
| *Candidatus Nitrosocosmicus franklandus* | 90 | Mb | 11.34297814 |
| *Candidatus Nitrosocosmicus franklandus* | 90 | Velifer | 10.99585623 |
| *Candidatus Nitrosocosmicus oleophilus* | -1 | Control | 6.3658671 |
| *Candidatus Nitrosocosmicus oleophilus* | -1 | Mb | 7.495205166 |
| *Candidatus Nitrosocosmicus oleophilus* | -1 | Velifer | 4.78111267 |
| *Candidatus Nitrosocosmicus oleophilus* | 0 | Mb | 6.316807782 |
| *Candidatus Nitrosocosmicus oleophilus* | 0 | Velifer | 4.916601482 |
| *Candidatus Nitrosocosmicus oleophilus* | 30 | Control | 7.810629851 |
| *Candidatus Nitrosocosmicus oleophilus* | 30 | Mb | 8.170391353 |
| *Candidatus Nitrosocosmicus oleophilus* | 30 | Velifer | 6.329043687 |
| *Candidatus Nitrosocosmicus oleophilus* | 90 | Control | 8.425832794 |
| *Candidatus Nitrosocosmicus oleophilus* | 90 | Mb | 7.815300408 |
| *Candidatus Nitrosocosmicus oleophilus* | 90 | Velifer | 7.560007647 |
| *Candidatus Nitrosopumilus koreensis* | -1 | Control | 1.322673118 |
| *Candidatus Nitrosopumilus koreensis* | -1 | Mb | 3.177918879 |
| *Candidatus Nitrosopumilus koreensis* | -1 | Velifer | 6.150567069 |
| *Candidatus Nitrosopumilus koreensis* | 0 | Mb | 2.089080378 |
| *Candidatus Nitrosopumilus koreensis* | 0 | Velifer | 5.682075219 |
| *Candidatus Nitrosopumilus koreensis* | 30 | Control | 1.039199225 |
| *Candidatus Nitrosopumilus koreensis* | 30 | Mb | 2.961714619 |
| *Candidatus Nitrosopumilus koreensis* | 30 | Velifer | 2.364032109 |
| *Candidatus Nitrosopumilus koreensis* | 90 | Control | 0.77590621 |
| *Candidatus Nitrosopumilus koreensis* | 90 | Mb | 1.490808709 |
| *Candidatus Nitrosopumilus koreensis* | 90 | Velifer | 1.880760013 |
| *Candidatus Nitrosopumilus sediminis* | -1 | Control | 0.639553776 |
| *Candidatus Nitrosopumilus sediminis* | -1 | Mb | 2.118108022 |
| *Candidatus Nitrosopumilus sediminis* | -1 | Velifer | 6.501107252 |
| *Candidatus Nitrosopumilus sediminis* | 0 | Mb | 1.560660293 |
| *Candidatus Nitrosopumilus sediminis* | 0 | Velifer | 5.413772401 |
| *Candidatus Nitrosopumilus sediminis* | 30 | Control | 0.47977902 |
| *Candidatus Nitrosopumilus sediminis* | 30 | Mb | 1.970825657 |
| *Candidatus Nitrosopumilus sediminis* | 30 | Velifer | 2.205122016 |
| *Candidatus Nitrosopumilus sediminis* | 90 | Control | 0.330530892 |
| *Candidatus Nitrosopumilus sediminis* | 90 | Mb | 0.849263651 |
| *Candidatus Nitrosopumilus sediminis* | 90 | Velifer | 1.730281479 |
| *Candidatus Nitrosopumilus* sp. *SW* | -1 | Control | 0.838468081 |
| *Candidatus Nitrosopumilus* sp. *SW* | -1 | Mb | 2.023264184 |
| *Candidatus Nitrosopumilus* sp. *SW* | -1 | Velifer | 4.587282346 |
| *Candidatus Nitrosopumilus* sp. *SW* | 0 | Mb | 1.438838332 |
| *Candidatus Nitrosopumilus* sp. *SW* | 0 | Velifer | 4.187349454 |
| *Candidatus Nitrosopumilus* sp. *SW* | 30 | Control | 0.739092546 |
| *Candidatus Nitrosopumilus* sp. *SW* | 30 | Mb | 1.921061682 |
| *Candidatus Nitrosopumilus* sp. *SW* | 30 | Velifer | 2.22763666 |
| *Candidatus Nitrosopumilus* sp. *SW* | 90 | Control | 0.405808285 |
| *Candidatus Nitrosopumilus* sp*. SW* | 90 | Mb | 1.032704806 |
| *Candidatus Nitrosopumilus* sp. *SW* | 90 | Velifer | 1.44107511 |
| *Candidatus Nitrososphaera evergladensis* | -1 | Control | 8.17983486 |
| *Candidatus Nitrososphaera evergladensis* | -1 | Mb | 7.616890358 |
| *Candidatus Nitrososphaera evergladensis* | -1 | Velifer | 6.472401688 |
| *Candidatus Nitrososphaera evergladensis* | 0 | Mb | 6.106427613 |
| *Candidatus Nitrososphaera evergladensis* | 0 | Velifer | 6.055544676 |
| *Candidatus Nitrososphaera evergladensis* | 30 | Control | 6.553421518 |
| *Candidatus Nitrososphaera evergladensis* | 30 | Mb | 7.45878394 |
| *Candidatus Nitrososphaera evergladensis* | 30 | Velifer | 8.813018029 |
| *Candidatus Nitrososphaera evergladensis* | 90 | Control | 3.228991357 |
| *Candidatus Nitrososphaera evergladensis* | 90 | Mb | 5.23200412 |
| *Candidatus Nitrososphaera evergladensis* | 90 | Velifer | 4.298270625 |
| *Candidatus Nitrososphaera gargensis* | -1 | Control | 31.38774792 |
| *Candidatus Nitrososphaera gargensis* | -1 | Mb | 24.61098344 |
| *Candidatus Nitrososphaera gargensis* | -1 | Velifer | 23.71144152 |
| *Candidatus Nitrososphaera gargensis* | 0 | Mb | 26.18168534 |
| *Candidatus Nitrososphaera gargensis* | 0 | Velifer | 24.06430334 |
| *Candidatus Nitrososphaera gargensis* | 30 | Control | 22.97264777 |
| *Candidatus Nitrososphaera gargensis* | 30 | Mb | 23.54335485 |
| *Candidatus Nitrososphaera gargensis* | 30 | Velifer | 32.0746606 |
| *Candidatus Nitrososphaera gargensis* | 90 | Control | 12.37573449 |
| *Candidatus Nitrososphaera gargensis* | 90 | Mb | 17.93653655 |
| *Candidatus Nitrososphaera gargensis* | 90 | Velifer | 14.7232401 |
| *Nitrosopumilus adriaticus* | -1 | Control | 0.495845217 |
| *Nitrosopumilus adriaticus* | -1 | Mb | 2.521248822 |
| *Nitrosopumilus adriaticus* | -1 | Velifer | 6.848037715 |
| *Nitrosopumilus adriaticus* | 0 | Mb | 1.682179216 |
| *Nitrosopumilus adriaticus* | 0 | Velifer | 5.999958344 |
| *Nitrosopumilus adriaticus* | 30 | Control | 0.468780348 |
| *Nitrosopumilus adriaticus* | 30 | Mb | 2.373740392 |
| *Nitrosopumilus adriaticus* | 30 | Velifer | 2.595784923 |
| *Nitrosopumilus adriaticus* | 90 | Control | 0.206430649 |
| *Nitrosopumilus adriaticus* | 90 | Mb | 0.744202796 |
| *Nitrosopumilus adriaticus* | 90 | Velifer | 1.91329749 |
| *Nitrososphaera viennensis* | -1 | Control | 7.811346638 |
| *Nitrososphaera viennensis* | -1 | Mb | 6.145017674 |
| *Nitrososphaera viennensis* | -1 | Velifer | 5.308452666 |
| *Nitrososphaera viennensis* | 0 | Mb | 5.228480487 |
| *Nitrososphaera viennensis* | 0 | Velifer | 5.064788492 |
| *Nitrososphaera viennensis* | 30 | Control | 4.709979443 |
| *Nitrososphaera viennensis* | 30 | Mb | 5.088777619 |
| *Nitrososphaera viennensis* | 30 | Velifer | 5.566030846 |
| *Nitrososphaera viennensis* | 90 | Control | 2.271710093 |
| *Nitrososphaera viennensis* | 90 | Mb | 3.774816184 |
| *Nitrososphaera viennensis* | 90 | Velifer | 2.787022725 |

Time: -1, 1 day before treatment application; 0, day of treatment application; 30 and 90, 30 and 90 days post-application.

**Supplementary Table 11 | Welch t-test analysis between time points**

| **Group 1** | **Group 2** | ***p*-value** | ***p*_adj_** |
| --- | --- | --- | --- |
| Control | Mb | 0.821 | 1 |
| Control | Velifer | 0.0113 | 0.034 |
| Mb | Velifer | 0.0144 | 0.0432 |

| **Comparison** | **t-statistic** | ***p*-value** | **FDR** | **DF** | **Mean ± SD Group 1** | **Mean ± SD Group 2** |
| --- | --- | --- | --- | --- | --- | --- |
| -1 vs 90 | 4.25 | 0.00 | 6.36E-04 | 44.97 | 6.783 ± 0.071 | 6.590 ± 0.282 |
| -1 vs 30 | 3.69 | 0.00 | 1.85E-03 | 43.58 | 6.783 ± 0.071 | 6.674 ± 0.160 |
| 0 vs 90 | 3.37 | 0.00 | 2.78E-03 | 53.63 | 6.753 ± 0.093 | 6.590 ± 0.282 |
| 0 vs 30 | 2.33 | 0.02 | 3.54E-02 | 53.53 | 6.753 ± 0.093 | 6.674 ± 0.160 |
| 30 vs 90 | 1.62 | 0.11 | 1.32E-01 | 65.10 | 6.674 ± 0.160 | 6.590 ± 0.282 |
| 0 vs -1 | -1.33 | 0.19 | 1.92E-01 | 34.27 | 6.753 ± 0.093 | 6.783 0.071 |

**Supplementary Table 12 | Control treatment metrics**

| **Species** | **Phylum** | **Abundance** | **Degree** | **Betweenness** | **Closeness** |
| --- | --- | --- | --- | --- | --- |
| *Pseudobacter ginsenosidimutans* | *Bacteroidetes* | 27.826036 | 14 | 51.07828283 | 0.04 |
| *Niabella ginsenosidivorans* | *Bacteroidetes* | 38.70213868 | 11 | 22.51313131 | 0.037037037 |
| *Paraflavitalea soli* | *Bacteroidetes* | 51.07216195 | 11 | 15.01313131 | 0.037037037 |
| *Flavisolibacter* sp. *17J28-1* | *Bacteroidetes* | 26.00703378 | 11 | 26.39393939 | 0.035714286 |
| *Niastella koreensis* | *Bacteroidetes* | 46.51015914 | 10 | 4.513131313 | 0.035714286 |
| *Flavisolibacter ginsenosidimutans* | *Bacteroidetes* | 29.92140294 | 10 | 4.513131313 | 0.035714286 |
| *Filimonas lacunae* | *Bacteroidetes* | 12.84631283 | 10 | 4.513131313 | 0.035714286 |
| *Flavisolibacter tropicus* | *Bacteroidetes* | 29.39078397 | 9 | 8.378787879 | 0.033333333 |
| *Niabella soli* | *Bacteroidetes* | 39.38210038 | 7 | 0 | 0.03125 |
| *Arachidicoccus ginsenosidivorans* | *Bacteroidetes* | 4.535863312 | 7 | 18 | 0.027027027 |
| *Arachidicoccus* sp. *BS20* | *Bacteroidetes* | 6.509145642 | 7 | 0 | 0.03030303 |
| *Chitinophaga pinensis* | *Bacteroidetes* | 14.28028207 | 5 | 1.055555556 | 0.026315789 |
| *Chitinophaga* sp. *XS-30* | *Bacteroidetes* | 9.061809724 | 5 | 0.972222222 | 0.026315789 |
| *Chitinophaga* sp. *T22* | *Bacteroidetes* | 4.534034666 | 5 | 3.055555556 | 0.026315789 |
| *Luteitalea pratensis* | *Acidobacteria* | 45.89209231 | 4 | 17 | 0.076923077 |
| *Chitinophaga caeni* | *Bacteroidetes* | 5.522163192 | 4 | 0.5 | 0.025641026 |
| *Chitinophaga* sp. *MD30* | *Bacteroidetes* | 5.1335031 | 4 | 0.5 | 0.024390244 |
| *Anaerolinea thermophila* | *Chloroflexi* | 32.89994935 | 3 | 2 | 0.333333333 |
| *Candidatus Solibacter usitatus* | *Acidobacteria* | 16.05926559 | 3 | 1 | 0.058823529 |
| *Nocardioides euryhalodurans* | *Actinobacteria* | 7.26145437 | 3 | 3 | 0.333333333 |
| *Granulicella mallensis* | *Acidobacteria* | 6.320788234 | 3 | 7 | 0.058823529 |
| *Gemmatimonas aurantiaca* | *Gemmatimonadetes* | 22.09725614 | 3 | 17 | 0.071428571 |
| *Gemmatirosa kalamazoonesis* | *Gemmatimonadetes* | 120.4697441 | 2 | 7 | 0.052631579 |
| *Nitrospira moscoviensis* | *Nitrospirae* | 51.94766724 | 2 | 1 | 0.5 |
| *Panacibacter ginsenosidivorans* | *Bacteroidetes* | 69.23934342 | 2 | 0 | 0.022727273 |
| *Pleomorphomonas* sp. *SM30* | *Proteobacteria* | 10.7121534 | 2 | 1 | 0.5 |
| *Candidatus Nitrosocosmicus franklandus* | *Thaumarchaeota* | 36.03310781 | 2 | 1 | 0.5 |
| *Egicoccus halophilus* | *Actinobacteria* | 13.17298094 | 2 | 1 | 0.5 |
| *Pelolinea submarina* | *Chloroflexi* | 25.38536862 | 2 | 0 | 0.25 |
| *Mucilaginibacter ginsenosidivorans* | *Bacteroidetes* | 4.832434724 | 2 | 0 | 0.024390244 |
| *Brevefilum fermentans* | *Chloroflexi* | 8.954681961 | 2 | 0 | 0.25 |
| *Candidatus Koribacter versatilis* | *Acidobacteria* | 8.758493696 | 2 | 0 | 0.052631579 |
| *Rubrobacter xylanophilus* | *Actinobacteria* | 38.61185925 | 1 | 0 | 1 |
| *Hyphomicrobium denitrificans* | *Proteobacteria* | 44.16428422 | 1 | 0 | 1 |
| *Streptomyces albogriseolus* | *Actinobacteria* | 21.14565012 | 1 | 0 | 1 |
| *Euzebya* sp. *DY32-46* | *Actinobacteria* | 13.52209725 | 1 | 0 | 0.333333333 |
| *Chelatococcus* sp. *CO-6* | *Proteobacteria* | 13.82923257 | 1 | 0 | 0.333333333 |
| *Candidatus Nitrosocosmicus exaquare* | *Thaumarchaeota* | 168.7959654 | 1 | 0 | 0.333333333 |
| *Nocardioides* sp. *JS614* | *Actinobacteria* | 7.960659753 | 1 | 0 | 0.2 |
| *Sphaerobacter thermophilus* | *Chloroflexi* | 13.4927056 | 1 | 0 | 1 |
| *Candidatus Promineofilum breve* | *Chloroflexi* | 91.70502735 | 1 | 0 | 1 |
| *Sorangium cellulosum* | *Proteobacteria* | 26.800277 | 1 | 0 | 1 |
| *Sphingomonas* sp*. AE3* | *Proteobacteria* | 19.914323 | 1 | 0 | 1 |
| *Rubrobacter radiotolerans* | *Actinobacteria* | 7.784013901 | 1 | 0 | 1 |
| *Hyphomicrobium* sp*. MC1* | *Proteobacteria* | 12.3708985 | 1 | 0 | 1 |
| *Nitrospira defluvii* | *Nitrospirae* | 10.58784763 | 1 | 0 | 0.333333333 |
| *Nitratireductor* sp*. SY7* | *Proteobacteria* | 5.137470109 | 1 | 0 | 0.333333333 |
| *Thermobifida fusca* | *Actinobacteria* | 17.79284103 | 1 | 0 | 1 |
| *Candidatus Nitrosocosmicus oleophilus* | *Thaumarchaeota* | 25.26202248 | 1 | 0 | 0.333333333 |
| *Pimelobacter simplex* | *Actinobacteria* | 4.535614283 | 1 | 0 | 0.2 |
| *Ilumatobacter coccineus* | *Actinobacteria* | 11.73454425 | 1 | 0 | 0.333333333 |
| *Pedobacter heparinus* | *Bacteroidetes* | 4.456148933 | 1 | 0 | 0.022222222 |
| *Gemmatimonas phototrophica* | *Gemmatimonadetes* | 11.19759839 | 1 | 0 | 0.038461538 |
| *Arachidicoccus* sp. *KIS59-12* | *Bacteroidetes* | 4.403817908 | 1 | 0 | 0.018181818 |
| *Nocardioides* sp*. 603* | *Actinobacteria* | 5.878720334 | 1 | 0 | 0.2 |
| *Skermanella pratensis* | *Proteobacteria* | 4.971995179 | 1 | 0 | 1 |
| *Candidatus Nitrospira inopinata* | *Nitrospirae* | 28.34465562 | 1 | 0 | 0.333333333 |
| *Caldilinea aerophila* | *Chloroflexi* | 18.41180639 | 1 | 0 | 1 |
| *Sandaracinus amylolyticus* | *Proteobacteria* | 10.44109293 | 1 | 0 | 1 |
| *Sphingomonas indica* | *Proteobacteria* | 8.15741666 | 1 | 0 | 1 |
| *Acidobacterium capsulatum* | *Acidobacteria* | 5.002831936 | 1 | 0 | 0.041666667 |
| *Roseiflexus castenholzii* | *Chloroflexi* | 5.993319717 | 1 | 0 | 0.2 |
| *Anaeromyxobacter* sp. *Fw109-5* | *Proteobacteria* | 7.218209915 | 1 | 0 | 0.047619048 |

**Supplementary Table 13 | Mb treatment network metrics**

| **Species** | **Phylum** | **Abundance** | **Degree** | **Betweenness** | **Closeness** |
| --- | --- | --- | --- | --- | --- |
| *Niastella koreensis* | *Bacteroidetes* | 45.55622025 | 9 | 10.5 | 0.066667 |
| *Flavisolibacter ginsenosidimutans* | *Bacteroidetes* | 27.88327814 | 9 | 5.866666667 | 0.066667 |
| *Paraflavitalea soli* | *Bacteroidetes* | 46.38433722 | 8 | 12.86666667 | 0.0625 |
| *Niabella ginsenosidivorans* | *Bacteroidetes* | 35.1328544 | 8 | 3.45 | 0.0625 |
| *Niabella soli* | *Bacteroidetes* | 37.93552378 | 7 | 1.95 | 0.055556 |
| *Pseudobacter ginsenosidimutans* | *Bacteroidetes* | 27.2648795 | 7 | 3.7 | 0.055556 |
| *Anaerolinea thermophila* | *Chloroflexi* | 29.71520505 | 6 | 1.666666667 | 0.125 |
| *Pelolinea submarina* | *Chloroflexi* | 22.55202467 | 6 | 1.666666667 | 0.125 |
| *Panacibacter ginsenosidivorans* | *Bacteroidetes* | 55.69499351 | 6 | 11.5 | 0.055556 |
| *Chloroflexus aurantiacus* | *Chloroflexi* | 5.504151604 | 6 | 6.666666667 | 0.125 |
| *Flavisolibacter tropicus* | *Bacteroidetes* | 26.01338782 | 5 | 0.916666667 | 0.05 |
| *Bradyrhizobium lablabi* | *Proteobacteria* | 5.505063819 | 5 | 57 | 0.02 |
| *Sorangium cellulosum* | *Proteobacteria* | 21.62911559 | 5 | 76.5 | 0.02439 |
| *Caldilinea aerophila* | *Chloroflexi* | 13.84559371 | 5 | 0.5 | 0.111111 |
| *Brevefilum fermentans* | *Chloroflexi* | 7.471558321 | 5 | 0.5 | 0.111111 |
| *Flavisolibacter* sp. *17J28-1* | *Bacteroidetes* | 26.37311386 | 5 | 0 | 0.05 |
| *Candidatus Promineofilum breve* | *Chloroflexi* | 57.55244886 | 4 | 0 | 0.090909 |
| *Planctomyces* sp*. SH-PL62* | *Planctomycetes* | 5.560502877 | 4 | 42.5 | 0.02 |
| *Blastochloris tepida* | *Proteobacteria* | 8.540433885 | 4 | 31 | 0.022222 |
| *Filimonas lacunae* | *Bacteroidetes* | 12.47745052 | 4 | 0.25 | 0.045455 |
| *Luteitalea pratensis* | *Acidobacteria* | 39.56944455 | 3 | 2 | 0.019231 |
| *Rhodopseudomonas palustris* | *Proteobacteria* | 18.48558155 | 3 | 72 | 0.025641 |
| *Rhodoplanes* sp*. Z2-YC6860* | *Proteobacteria* | 17.23545246 | 3 | 60 | 0.02381 |
| *Anaeromyxobacter dehalogenans* | *Proteobacteria* | 8.41308516 | 3 | 0.5 | 0.018182 |
| *Stackebrandtia nassauensis* | *Actinobacteria* | 7.565956197 | 3 | 2 | 0.333333 |
| *Roseiflexus castenholzii* | *Chloroflexi* | 4.726387907 | 3 | 0 | 0.090909 |
| *Brevundimonas naejangsanensis* | *Proteobacteria* | 3.965230339 | 3 | 4 | 0.2 |
| *Candidatus Nitrosocosmicus franklandus* | *Thaumarchaeota* | 28.63488818 | 2 | 1 | 0.5 |
| *Rubrobacter xylanophilus* | *Actinobacteria* | 53.3951744 | 2 | 1 | 0.5 |
| *Caulobacter segnis* | *Proteobacteria* | 5.333508737 | 2 | 0 | 0.142857 |
| *Gemmatirosa kalamazoonesis* | *Gemmatimonadetes* | 85.86087493 | 2 | 1 | 0.5 |
| *Sphingomonas* sp*. AE3* | *Proteobacteria* | 18.55963617 | 2 | 1 | 0.5 |
| *Candidatus Nitrososphaera evergladensis* | *Thaumarchaeota* | 15.60481278 | 2 | 1 | 0.5 |
| *Caulobacter vibrioides* | *Proteobacteria* | 6.485225375 | 2 | 3 | 0.166667 |
| *Pseudolabrys* sp*. FHR47* | *Proteobacteria* | 8.288002332 | 2 | 0 | 0.015385 |
| *Hyphomonas* sp*. CACIAM 19H1* | *Proteobacteria* | 5.153462689 | 2 | 0 | 0.142857 |
| *Streptomyces albogriseolus* | *Actinobacteria* | 21.23017992 | 2 | 0 | 0.25 |
| *Nocardioides euryhalodurans* | *Actinobacteria* | 6.258717658 | 2 | 1 | 0.5 |
| *Candidatus Solibacter usitatus* | *Acidobacteria* | 12.33963568 | 2 | 0.5 | 0.012821 |
| *Starkeya novella* | *Proteobacteria* | 6.458163929 | 2 | 0 | 0.015385 |
| *Thermobifida fusca* | *Actinobacteria* | 16.59219414 | 2 | 0 | 0.25 |
| *Chitinophaga pinensis* | *Bacteroidetes* | 12.62176419 | 2 | 0 | 0.041667 |
| *Anaeromyxobacter* sp*. Fw109-5* | *Proteobacteria* | 5.450264024 | 2 | 0 | 0.017857 |
| *Anaeromyxobacter* sp. *K* | *Proteobacteria* | 4.05056867 | 2 | 7.5 | 0.015625 |
| *Granulicella mallensis* | *Acidobacteria* | 5.421634614 | 2 | 7.5 | 0.015625 |
| *Hyphomicrobium denitrificans* | *Proteobacteria* | 21.8135486 | 1 | 0 | 1 |
| *Nitrospira moscoviensis* | *Nitrospirae* | 58.98251781 | 1 | 0 | 1 |
| *Candidatus Nitrosocosmicus exaquare* | *Thaumarchaeota* | 119.4931507 | 1 | 0 | 0.333333 |
| *Candidatus Nitrososphaera gargensis* | *Thaumarchaeota* | 53.64203944 | 1 | 0 | 0.333333 |
| *Phenylobacterium zucineum* | *Proteobacteria* | 7.488633156 | 1 | 0 | 0.111111 |
| *Nocardioides* sp. *JS614* | *Actinobacteria* | 6.610539422 | 1 | 0 | 0.333333 |
| *Pseudolabrys taiwanensis* | *Proteobacteria* | 11.21648424 | 1 | 0 | 0.016393 |
| *Candidatus Nitrosocosmicus oleophilus* | *Thaumarchaeota* | 19.98156257 | 1 | 0 | 0.333333 |
| *Hyphomicrobium* sp*. MC1* | *Proteobacteria* | 6.642464899 | 1 | 0 | 1 |
| *Rubrobacter radiotolerans* | *Actinobacteria* | 10.64172515 | 1 | 0 | 0.333333 |
| *Nitrospira defluvii* | *Nitrospirae* | 11.90608181 | 1 | 0 | 1 |
| *Bradyrhizobium icense* | *Proteobacteria* | 4.661050996 | 1 | 0 | 0.015152 |
| *Roseiflexus* sp*. RS-1* | *Chloroflexi* | 3.848094827 | 1 | 0 | 0.071429 |
| *Gemmatimonas phototrophica* | *Gemmatimonadetes* | 7.11417469 | 1 | 0 | 0.333333 |
| *Arachidicoccus* sp*. BS20* | *Bacteroidetes* | 6.384355544 | 1 | 0 | 0.034483 |
| *Sphingomonas indica* | *Proteobacteria* | 7.133778604 | 1 | 0 | 0.333333 |
| *Nitrososphaera viennensis* | *Thaumarchaeota* | 11.47439106 | 1 | 0 | 0.333333 |
| *Mucilaginibacter ginsenosidivorans* | *Bacteroidetes* | 4.440425184 | 1 | 0 | 0.037037 |
| *Nocardioides* sp. *SB3-45* | *Actinobacteria* | 5.397406738 | 1 | 0 | 0.333333 |
| *Rubrobacter indicoceani* | *Actinobacteria* | 5.18731629 | 1 | 0 | 0.333333 |
| *Bradyrhizobium diazoefficiens* | *Proteobacteria* | 5.239240112 | 1 | 0 | 0.016393 |
| *Nocardiopsis dassonvillei* | *Actinobacteria* | 5.934862644 | 1 | 0 | 0.2 |
| *Gemmatimonas aurantiaca* | *Gemmatimonadetes* | 17.12876293 | 1 | 0 | 0.333333 |
| *Sphingosinicella* sp. *BN140058* | *Proteobacteria* | 8.958711735 | 1 | 0 | 0.333333 |
| *Blastochloris viridis* | *Proteobacteria* | 4.936674163 | 1 | 0 | 0.015152 |

**Supplementary Table 14 | Velifer treatment network metrics**

| **Species** | **Phylum** | **Abundance** | **Degree** | **Betweenness** | **Closeness** |
| --- | --- | --- | --- | --- | --- |
| *Niabella soli* | *Bacteroidetes* | 49.10546804 | 12 | 14.56423926 | 0.043478261 |
| *Chitinophaga pinensis* | *Bacteroidetes* | 16.62502183 | 11 | 11.84367336 | 0.041666667 |
| *Paraflavitalea soli* | *Bacteroidetes* | 64.89450667 | 10 | 16.05054812 | 0.041666667 |
| *Niabella ginsenosidivorans* | *Bacteroidetes* | 49.41948815 | 10 | 23.58032861 | 0.041666667 |
| *Niastella koreensis* | *Bacteroidetes* | 62.7155684 | 10 | 4.237957212 | 0.04 |
| *Flavisolibacter tropicus* | *Bacteroidetes* | 39.45138595 | 10 | 21.22056591 | 0.04 |
| *Filimonas lacunae* | *Bacteroidetes* | 16.42209399 | 10 | 4.237957212 | 0.04 |
| *Flavisolibacter ginsenosidimutans* | *Bacteroidetes* | 35.2983919 | 9 | 6.996107395 | 0.04 |
| *Flavisolibacter* sp. *17J28-1* | *Bacteroidetes* | 36.60519783 | 9 | 4.582908637 | 0.035714286 |
| *Pseudobacter ginsenosidimutans* | *Bacteroidetes* | 41.16919927 | 6 | 0.685714286 | 0.03030303 |
| *Arachidicoccus ginsenosidivorans* | *Bacteroidetes* | 5.868712556 | 6 | 10.58833148 | 0.03030303 |
| *Chitinophaga caeni* | *Bacteroidetes* | 7.662330005 | 6 | 0.166666667 | 0.029411765 |
| *Candidatus Nitrosopumilus koreensis* | *Thaumarchaeota* | 8.130193105 | 5 | 0 | 0.2 |
| *Panacibacter ginsenosidivorans* | *Bacteroidetes* | 83.67552755 | 5 | 6.245001858 | 0.029411765 |
| *Candidatus Nitrosopumilus* sp. *SW* | *Thaumarchaeota* | 6.190041656 | 5 | 0 | 0.2 |
| *Candidatus Nitrosopumilus sediminis* | *Thaumarchaeota* | 7.912069763 | 5 | 0 | 0.2 |
| *Nitrosopumilus adriaticus* | *Thaumarchaeota* | 8.727355756 | 5 | 0 | 0.2 |
| *Nitrosopumilus piranensis* | *Thaumarchaeota* | 5.300384934 | 5 | 0 | 0.2 |
| *Nitrosopumilus maritimus* | *Thaumarchaeota* | 5.054449712 | 5 | 0 | 0.2 |
| *Nitrospira moscoviensis* | *Nitrospirae* | 64.30370827 | 3 | 0.5 | 0.333333333 |
| *Gemmatirosa kalamazoonesis* | *Gemmatimonadetes* | 83.59321698 | 3 | 6 | 0.142857143 |
| *Luteitalea pratensis* | *Acidobacteria* | 40.30198946 | 3 | 7 | 0.142857143 |
| *Nitrospira defluvii* | *Nitrospirae* | 12.84196172 | 3 | 0.5 | 0.333333333 |
| *Arachidicoccus* sp. *KIS59-12* | *Bacteroidetes* | 5.550677949 | 3 | 0 | 0.027027027 |
| *Chitinophaga* sp*. T22* | *Bacteroidetes* | 6.092288674 | 3 | 0 | 0.027027027 |
| *Arachidicoccus* sp*. BS20* | *Bacteroidetes* | 7.795143286 | 3 | 0 | 0.027777778 |
| *Rubrobacter xylanophilus* | *Actinobacteria* | 66.41549607 | 2 | 0 | 0.5 |
| *Nitrospira japonica* | *Nitrospirae* | 30.91050578 | 2 | 0 | 0.25 |
| *Rubrobacter radiotolerans* | *Actinobacteria* | 13.04499949 | 2 | 0 | 0.5 |
| *Candidatus Nitrospira inopinata* | *Nitrospirae* | 45.63234374 | 2 | 0 | 0.25 |
| *Gemmatimonas aurantiaca* | *Gemmatimonadetes* | 17.54642789 | 2 | 0 | 0.1 |
| *Rubrobacter indicoceani* | *Actinobacteria* | 6.187629892 | 2 | 0 | 0.5 |
| *Gemmatimonas phototrophica* | *Gemmatimonadetes* | 9.162968085 | 2 | 0 | 0.1 |
| *Pedobacter ginsengisoli* | *Bacteroidetes* | 4.346900701 | 2 | 0 | 0.020833333 |
| *Brevefilum fermentans* | *Chloroflexi* | 5.853131494 | 2 | 1 | 0.5 |
| *Candidatus Nitrosocosmicus exaquare* | *Thaumarchaeota* | 118.6534967 | 1 | 0 | 1 |
| *Streptomyces albogriseolus* | *Actinobacteria* | 22.94857827 | 1 | 0 | 1 |
| *Hyphomicrobium denitrificans* | *Proteobacteria* | 21.75832823 | 1 | 0 | 1 |
| *Pseudomonas* sp. *PE08* | *Proteobacteria* | 11.43393249 | 1 | 0 | 1 |
| *Anaerolinea thermophila* | *Chloroflexi* | 26.05847037 | 1 | 0 | 0.333333333 |
| *Candidatus Promineofilum breve* | *Chloroflexi* | 87.03222203 | 1 | 0 | 1 |
| *Gramella* sp. *MAR_2010_147* | *Bacteroidetes* | 7.03275465 | 1 | 0 | 1 |
| *Pelolinea submarina* | *Chloroflexi* | 19.82341333 | 1 | 0 | 0.333333333 |
| *Candidatus Nitrosocosmicus franklandus* | *Thaumarchaeota* | 25.33308778 | 1 | 0 | 1 |
| *Thermobifida fusca* | *Actinobacteria* | 16.87320392 | 1 | 0 | 1 |
| *Hyphomicrobium* sp. *MC1* | *Proteobacteria* | 6.649495248 | 1 | 0 | 1 |
| *Pseudomonas* sp*. TCU-HL1* | *Proteobacteria* | 5.561421275 | 1 | 0 | 1 |
| *Caldilinea aerophila* | *Chloroflexi* | 14.34401869 | 1 | 0 | 1 |
| *Candidatus Solibacter usitatus* | *Acidobacteria* | 15.78128718 | 1 | 0 | 0.090909091 |
| *Arachidicoccus* sp*. B3-10* | *Bacteroidetes* | 5.218144529 | 1 | 0 | 0.024390244 |
| *Chloroflexus aurantiacus* | *Chloroflexi* | 5.147659636 | 1 | 0 | 0.090909091 |
| *Gramella* sp. *MAR_2010_102* | *Bacteroidetes* | 4.966465971 | 1 | 0 | 1 |

**Supplementary Table 1****5 | Logfold change of statistically significant species in Velifer and Mb treatments compared to the control at 90 days post-application**

| **ID** | **logFC (90_control/90_Mb)** | ***p*_adj_** | **logFC (90_control/90_velifer)** | ***p*_adj_** |
| --- | --- | --- | --- | --- |
| *Pseudomonas_furukawaii* | -1.684745909 | 1.14E-02 | -3.005641373 | 1.72E-19 |
| *Pseudomonas_*sp.*_PE08* | -1.809199198 | 0.002036 | -3.177183311 | 6E-16 |
| *Azotobacter_chroococcum* | -1.06276721 | 0.01679 | -3.79472023 | 1.11E-09 |
| *Pseudomonas_*sp*._TCU-HL1* |  |  | -2.775794009 | 1.02E-08 |
| *Pseudomonas_mendocina* | -1.462968193 | 0.049803 | -1.755888908 | 4.63E-07 |
| *Pseudobacter_ginsenosidimutans* |  |  | -1.508251768 | 8.66E-07 |
| *Azotobacter_salinestris* |  |  | -3.268480535 | 2.8E-06 |
| *Niastella_koreensis* |  |  | -1.337303359 | 5.47E-06 |
| *Pseudomonas_*sp.*_DY-1* |  |  | -2.265698804 | 5.47E-06 |
| *Stackebrandtia_nassauensis* |  |  | -2.926679253 | 1.14E-05 |
| *Paraflavitalea_soli* |  |  | -1.281420153 | 2.45E-05 |
| *Pseudomonas_resinovorans* |  |  | -2.819657768 | 3.58E-05 |
| *Niabella_ginsenosidivorans* |  |  | -1.183476138 | 1.23E-04 |
| *Flavisolibacter_*sp.*_17J28-1* |  |  | -1.361281243 | 1.26E-04 |
| *Pseudomonas_oleovorans* |  |  | -2.437788712 | 0.000167 |
| *Nitrosopumilus_adriaticus* |  |  | -2.452583411 | 0.000167 |
| *Flavisolibacter_tropicus* |  |  | -1.39459323 | 0.000167 |
| *Niabella_soli* |  |  | -1.017628714 | 1.67E-04 |
| *Pseudomonas_*sp*._LPH1* |  |  | -1.49489651 | 0.000318 |
| *Flavisolibacter_ginsenosidimutans* |  |  | -1.170074519 | 0.000399 |
| *Chitinophaga_*sp*._XS-30* |  |  | -1.262839061 | 0.000786 |
| *Panacibacter_ginsenosidivorans* |  |  | -1.15234728 | 0.000786 |
| *Filimonas_lacunae* |  |  | -1.245663868 | 0.001205 |
| *Candidatus_Nitrosopumilus_sediminis* |  |  | -1.849130262 | 0.001214 |
| *Amycolatopsis_albispora* |  |  | -1.330607141 | 0.001765 |
| *Hyphomicrobium_denitrificans* | 1.083991418 | 0.000113 | 1.065824128 | 2.11E-03 |
| *Woeseia_oceani* |  |  | 1.037476816 | 0.002267 |
| *Acinetobacter_chinensis* |  |  | -1.467503211 | 0.00245 |
| *Chitinophaga_*sp.*_T22* |  |  | -1.248996329 | 0.00271 |
| *Hyphomicrobium_*sp.*_MC1* | 1.017066803 | 0.00095 | 0.926792143 | 0.002899 |
| *Arachidicoccus_*sp.*_B3-10* |  |  | -1.468201025 | 0.004999 |
| *Thermothelomyces_thermophilus* |  |  | -1.180152343 | 0.006012 |
| *Blackfly_microvirus_SF02* | -5.192080159 | 0.002036 | -4.308115101 | 0.010683 |
| *Pseudomonas_alcaliphila* |  |  | -3.213546278 | 1.07E-02 |
| *Chitinophaga_caeni* |  |  | -1.209355785 | 0.010683 |
| *Candidatus_Nitrosopumilus_*sp.*_SW* |  |  | -1.316899937 | 0.011023 |
| *Arachidicoccus_*sp*._BS20* |  |  | -1.087453339 | 0.014849 |
| *Streptomyces_*sp*._ICC4* |  |  | -2.637567076 | 0.014849 |
| *Thioflavicoccus_mobilis* |  |  | 1.345281735 | 1.70E-02 |
| *Pseudomonas_*sp*._SWI7* |  |  | 2.758084532 | 4.85E-02 |
| *Streptomyces_armeniacus* |  |  | -1.669773058 | 0.048511 |
| *Anseongella_ginsenosidimutans* | -3.958745255 | 1.49E-05 |  |  |
| *Metarhizium_brunneum* | -2.752413693 | 0.016966 |  |  |
